# Evidence for hierarchical representations of written and spoken words from an open-science human neuroimaging dataset

**DOI:** 10.1101/2025.09.07.674741

**Authors:** Suneel Banerjee, Kimberly Jin, Plamen Nikolov, Philip Cho, Vishnu R. Pendri, Lillian Chang, Srikanth R. Damera, Maximilian Riesenhuber

## Abstract

Reading and speech recognition rely on multi-level processing that builds from basic visual or sound features to complete word representations, yet details of these processing hierarchies (in particular those for spoken words) are still poorly understood. We re-analyzed the functional magnetic resonance imaging (fMRI) data provided in the Mother Of all Unification Studies (MOUS) open-science dataset by using parametric regressions of word frequency and sublexical unit (bigram or syllable) frequency during reading and speech listening tasks in order to elucidate lexical processing hierarchies in the visual and auditory modalities. We first validated our approach in the written word domain, where the technique identified significant correlations for word frequency in the left mid-fusiform cortex (at the location of the Visual Word Form Area) with a left occipital region tracking bigram frequency, compatible with prior reports. During listening, low-frequency spoken words elicited greater responses in a left mid-superior temporal region consistent with the recently-described Auditory Word Form Area (AWFA), while a more posterior region of the superior temporal gyrus was sensitive to syllable frequency. Activation in the left inferior frontal gyrus correlated with both written and spoken word frequency. These findings demonstrate parallel hierarchical organizations in the anteroventral visual and auditory streams, with modality-specific lexica and upstream sublexical representations that converge in higher-order language areas.

## 1. Introduction

The neural substrates underlying reading and speech recognition are two remarkable instances of hierarchical processing in the human brain. Reading leverages the ventral visual (“what”) stream. This stream is thought to be organized in a posterior-to-anterior hierarchy of neural selectivity increasingly complex shape features (Hubel et al., 1977; Kravitz et al., 2013). Applied to orthography, this ‘simple-to-complex’ model predicts that increasingly complex sublexical representations (e.g., those of bigrams and trigrams) emerge in progressively anterior regions in the ventral occipital-temporal cortex (vOT) (Vinckier et al., 2007), culminating with full lexical representations in the Visual Word Form Area (VWFA) in the left mid-fusiform cortex(Cohen et al., 2002; Glezer et al., 2009).

Motivated by findings of lower responses to familiar vs. unfamiliar objects in fMRI (Chao et al., 2002), early studies tested the hypothesis that, similarly, brain areas containing lexical orthographic representations should respond less to familiar (i.e., high frequency) than to unfamiliar (i.e., low frequency) words. This prediction was confirmed in the VWFA (Kronbichler et al., 2004). In parallel, word frequency has been shown to modulate reaction times in lexical decision tasks in both sensory modalities (Goh et al., 2009; Keuleers, Brysbaert, et al., 2010; NEW et al., 2007) Subsequent neuroimaging and human electrocorticography (ECoG) studies have provided further evidence for the frequency effect for written words in the VWFA (Fiez et al., 1999; Graves et al., 2010; Hauk et al., 2008; Joubert et al., 2004; Woolnough et al., 2021). The existence of a ‘visual dictionary’ in the VWFA shaped by experience was also demonstrated through fMRI-rapid adaptation techniques (Glezer et al., 2009, 2015) and subsequently confirmed by ECoG (Glezer et al., 2015; Lochy et al., 2018; Woolnough et al., 2021).

Other studies have applied a similar approach to probe sublexical representations using bigram frequencies (Graves et al., 2010; Woolnough et al., 2021) based on the hypothesis that lifelong exposure to written language leads to adaptive learning processes that guide the tuning of feature detectors in the ventral visual stream for frequently-encountered fragments of words (e.g., “GH” vs “GZ” in English) (Vinckier et al., 2007). These studies found significant inverse correlations of blood oxygen level-dependent (BOLD) responses with bigram frequency in a region posterior to the VWFA (later labeled the “occipital word form area,” OWFA) (Strother et al., 2016), thus demonstrating how the class of orthographic stimuli instantiates the general ‘simple-to-complex’ hierarchical processing principle of the ventral visual stream (Riesenhuber & Poggio, 1999). Just as in the case of the VWFA, this finding of sublexical selectivity in the OWFA obtained with the frequency correlation approach was also validated and confirmed through other approaches using fMRI and ECoG (Glezer et al., 2015; Seghier et al., 2012; Strother et al., 2016; Woolnough et al., 2021).

In contrast to hierarchical (i.e., lexical and sublexical) representations for written words in the visual system, the representations of spoken words in the brain’s auditory system are much less well understood. As a general principle, the auditory system, like the visual system, is also thought to be organized into two streams in the cortex: The antero-ventral stream (aka the “what” stream), projects from primary auditory cortex in Heschl’s gyrus to the inferior frontal gyrus via the mid and anterior superior temporal gyrus and is specialized for identifying acoustic stimuli, including spoken words. This stream is likewise organized along a simple-to-complex hierarchy (Kell et al., 2018; Norman-Haignere et al., 2022; Rauschecker & Scott, 2009), wherein progressively anterior neuronal populations in the superior temporal cortex integrate over longer timescales, permitting the representation of increasingly complex auditory “objects.” Using an fMRI-rapid adaptation approach very similar to our earlier study of lexical representations in the VWFA (Glezer et al., 2009, 2015), we recently provided evidence of neural representations for individual spoken words in the mid-anterior superior temporal gyrus (STG) (Damera et al., 2023), the site of the putative “Auditory Word Form Area*”* (AWFA) (Cohen & Dehaene, 2004; DeWitt & Rauschecker, 2012), sharpening the parallels between the brain’s encoding schemes for written and spoken words

Despite its successes in elucidating hierarchical representations for written words, word and word-part frequency have not previously been used to probe lexical or sublexical representations in the auditory system. If the recently-elucidated AWFA stores representations of spoken words, we would hypothesize stronger responses to infrequent vs. frequent auditory words (i.e., an inverse correlation between BOLD response level and word frequency), in analogy to the response profile of the VWFA. In addition (in analogy with the neural representation of bigrams posterior to the VWFA in the ventral visual stream), we would predict the existence of an area showing a frequency effect for spoken word-parts, such as syllables, posterior to the AWFA (a “mid-anterior auditory word form area”, mAWFA, as it were), yet anterior to Heschl’s gyrus, which performs early acoustic processing. Confirming these hypotheses would show intriguing parallels between written and spoken word processing systems in the brain and further our understanding of hierarchical processing as a general principle across sensory modalities.

These are the predictions we tested in the present study. A notable feature of our approach was its use of the Mother-Of-all Unification Studies (MOUS) dataset, a large, open-science human neuroimaging dataset that contains fMRI data collected as subjects completed single-word reading and speech listening tasks (n = 204, 102 reading and 102 speech listening) (Schoffelen et al., 2019). We re-analyzed this dataset by employing parametric regression to identify brain regions whose activity was inversely related to the frequency of words and their sublexical components, both in the written and auditory domains. Specifically, we first validated our approach using the written word/reading dataset (which identified lexical selectivity in the VWFA and sublexical selectivity in the OWFA regions, in line with the literature), and then applied the same approach to probe the hierarchical representation of spoken words in the brain.

Our analyses revealed strong parallels between lexical processing hierarchies in the auditory and written domains, providing evidence for separate word recognition hierarchies for speech and reading as well as downstream convergence areas in the prefrontal cortex. We found that word frequency regressors localize the VWFA during reading tasks and the AWFA during speech listening. Sublexical frequency regressors (constructed from bigram frequency and syllable frequency in the visual and auditory cases respectively) revealed peaks posterior to these lexical regions, identifying sites of intermediate orthographic and phonetic selectivity, respectively. These results provide support for lexical representations in the newly-reported AWFA and syllable representations posterior to it in the STG, situating the AWFA at the top of a word-recognition hierarchy in the auditory “what” pathway and suggesting a role analogous to that of the VWFA in the visual ventral stream.

## 2. Methods

### 2.1. Dataset

We utilized the Mother-Of-all Unification Studies (MOUS), which contains fMRI (task and resting-state), DWI, and MEG data from 204 subjects (100 males, mean age 22, range 18-33 years). Text in this section is largely reproduced from Schoffelen et al., 2019, in which the dataset was first described. Half (n = 102) of the subjects completed a reading task which consisted of Dutch sentences and scrambled word lists presented one word at a time, while the other half completed a speech listening task that consisted of spoken versions of the same linguistic stimuli. The full stimulus set contained 360 sentences, from which 548 scrambled sentences were also derived for a total of 908 linguistic stimuli. Each subject was presented with 60 scrambled word lists (‘Woorden’) and 60 sentences (‘Zinnen’) while undergoing fMRI scanning. Subjects were not exposed to scrambled word lists corresponding to sentences they had been presented with. Word lists and sentences varied between 9 and 15 words in length. Sentences or word lists were presented word-by-word with a mean duration of 351 ms for each word (minimum of 300 ms and maximum of 1400 ms). Specifically, the visual presentation rate of the stimuli was determined in relation to the duration of the audio recording of spoken versions of the sentences and the word lists (‘audiodur’ variable in the MOUS dataset), taking into account both the number of letters (‘sumnletters’) and words (‘nwords’) in the whole sentence and the number of letters within each word (‘nletters’). The duration of a single word (in ms) was determined by the MOUS study team as:

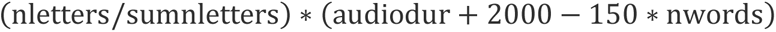

The auditory versions of the stimuli were recorded by a female native Dutch speaker. The word lists were pronounced with a neutral prosody and a clear pause in between each word. In order to check for compliance, 20% of the trials were followed by a ‘Yes’/‘No’ question about the content of the previous sentence/word list. Half of the questions on the sentences addressed the content of the sentence (e.g. *“Did grandma give a cookie to the girl*?”) whereas the other half, and all the questions about the word lists, addressed one of the main content words (e.g., “*Was the word ‘grandma’ mentioned?”*).

### 2.2. MRI Data Acquisition (Schoffelen et al., 2019)

The data were acquired with a SIEMENS Trio 3 T scanner using a 32-channel head coil. The collection order of the data analyzed here was as follows: 1) resting-state fMRI, 2) task-based fMRI, pause, 3) structural image.

#### 2.2.1. Task fMRI

T2*-weighted functional echo planar blood oxygenation level dependent (EPI-BOLD) data were acquired during the task. A single echo 2D ascending slice acquisition sequence (with partial brain coverage) was implemented with the following specifications: Volume TR = 2.00 s, TE = 35 ms, 90 degree flip-angle, 29 oblique slices (position: L.076 A6.4 H16.6, orientation: T > C-5.8 > S-1.2, Phase encoding direction: A >> P, rotation: −0.80 degrees), slice-matrix size (base resolution) = 64 × 64, slice thickness = 3.0 mm, slice gap 0.5 mm, FOV = 224 mm, voxel size (x = FOV_x_/N_x_, y = FOV_y_/N_y_, z = slice thickness, anisotropic voxel size = 3.5 × 3.5 × 3.0 mm).

#### 2.2.2. Structural imaging

A T1-weighted magnetization-prepared rapid gradient-echo (MP-RAGE) pulse sequence was used for the structural images, with the following parameters: volume TR = 2300 ms, TE = 3.03 ms, 8 degree flip-angle, 1 slab, slice-matrix size = 256 × 256, slice thickness = 1 mm, field of view = 256 mm, isotropic voxel-size = 1.0 × 1.0 × 1.0 mm. A vitamin-E capsule was placed as fiducial behind the right ear to allow a visual identification of left-right consistency.

#### 2.2.3. Anatomical data preprocessing (auto-generated)

A total of 200 T1-weighted (T1w) images were found within the input BIDS dataset. All of them were bias-field corrected and skull-stripped with ANTS 2.2.0 (Avants et al., 2008; Tustison et al., 2010). Brain tissue segmentation of cerebrospinal fluid (CSF), white-matter (WM) and gray-matter (GM) was performed on the brain-extracted T1w using FSL 5.0.9 (Bernal et al., 2021). A T1w-reference map was computed using FreeSurfer 6.0.1 (Reuter et al., 2010) Volume-based spatial normalization to two standard spaces (MNI152NLin6Asym and MNI152NLin2009cAsym) was performed with ANTs 2.2.0) (Chung et al., 2010; Evans et al., 2012).

#### 2.2.4. Functional data preprocessing (auto-generated)

Results in this manuscript come from data preprocessed with fMRIPrep 20.1.1 (Esteban et al., 2019), which is based on Nipype 1.5.0 (Gorgolewski et al., 2011). Due to missing or invalid anatomical (T1-weighted) or functional scans, 4 subjects (1 from the auditory group, A2091, and 3 from the visual group, V1050, V1109, V1071) were excluded from all further analyses. For each of the single BOLD runs found per subject (across all tasks and sessions), the following preprocessing was performed: First, a reference volume and its skull-stripped version were generated using fMRIPrep. Head-motion parameters with respect to the BOLD reference (transformation matrices, and six corresponding rotation and translation parameters) were estimated before any spatiotemporal filtering using FSL 5.0.9 (Jenkinson et al., 2002). BOLD runs were slice-time corrected using AFNI 20160207 (Cox & Hyde, 1997). Susceptibility distortion correction (SDC) was omitted. The BOLD reference was then co-registered to the T1w reference using FSL 5.0.9 (Greve & Fischl, 2009; Jenkinson & Smith, 2001). The BOLD time-series (including slice-timing correction) were resampled onto their original, native space by applying the transforms to correct for head-motion. The BOLD time-series were also resampled into standard space, generating a preprocessed BOLD run in MNI152NLin6Asym space. Functional volumes were resampled to isotropic 2 × 2 × 2 mm voxels and smoothed by 6mm (FWHM) in each direction. Several confounding time-series were calculated based on the preprocessed BOLD: framewise displacement (FD) (Jenkinson et al., 2002; Power et al., 2011), DVARS (D referring to temporal derivative of timecourses, VARS referring to RMS variance over voxels) and three region-wise global signals. FD and DVARS are calculated for each functional run (Power et al., 2011). The three global signals were extracted within the CSF, the WM, and the whole-brain masks. Additionally, a set of physiological regressors were extracted to allow for component-based noise correction (“CompCor”) (Behzadi et al., 2007). The head-motion estimates calculated in the correction step were also placed within the corresponding confounds file (Satterthwaite et al., 2013). Frames that exceeded a threshold of 0.4 mm FD or 1.5 standardized DVARS were annotated as motion outliers. Volumetric resamplings were performed using ANTs (Lanczos, 1964) and surface resamplings were performed using FreeSurfer. Many internal operations of fMRIPrep use Nilearn* 0.6.2, mostly within the functional processing workflow. For more details of the pipeline, see the section corresponding to workflows in fMRIPrep’s documentation (https://fmriprep.org/en/latest/workflows.html).

### 2.3. Word, bigram, and syllable frequency calculations

We sourced the frequencies of the Dutch words used in the fMRI tasks of the MOUS dataset from the SUBTLEX-NL Dutch language database (Keuleers, Brysbaert, et al., 2010). This database employs a text corpus derived from film subtitles, thus approximating everyday word usage statistics more accurately than other written corpora. To construct our word frequency regressors, we used the provided ‘Zipf’ frequency, which is calculated as follows:

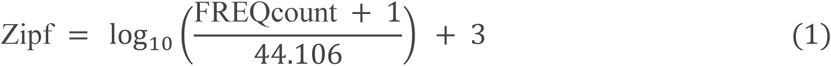

Where *FREQcount* represents the number of times a word appears in the SUBTLEX-NL text corpus. Log-transformed measures of word frequency have been shown to better explain variance in reaction time during lexical decision tasks than linear measures (Brysbaert et al., 2011, 2018; Keuleers, Diependaele, et al., 2010). Most studies of the word frequency effect have employed log-transformed word frequency values from SUBTLEX or CELEX lexical databases (Graves et al., 2010; Kronbichler et al., 2004; Woolnough et al., 2021).

We calculated individual bigram frequency by summing the FREQcounts of all words containing a certain two-letter combination, then taking the log of this value (Graves et al., 2010; Kronbichler et al., 2004).

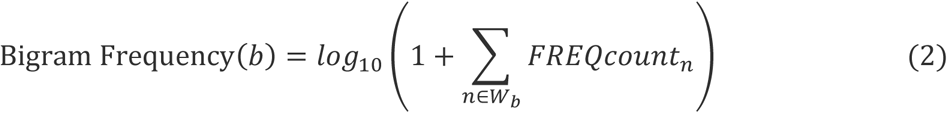

Where *b* is the bigram of interest (e.g., “th”, “an”), *W*_*b*_ is the set of word indices whose orthographic form contains *b*, and *FREQcount*_*n*_ is the value of FREQcount for the *n*^*th*^word. For each word that appeared in the fMRI tasks, we determined the mean, minimum and maximum bigram frequency found in a word.

To calculate syllable frequency, every native Dutch word found in the SUBTLEX-NL database was divided into its constituent syllables. We applied a novel, validated deep learning framework (Lathouwers et al., 2025), which slides a 5-letter window across each word, summarizes the local context with a convolutional neural network, then applies a bidirectional long short-term memory (LSTM) recurrent neural network followed by a conditional random field to decide, for every character, whether it begins a new syllable. Following the same process used to calculate bigram frequency (Equation 2), we then summed the frequencies (FREQcount) of all words containing each unique syllable and log-transformed this sum (base 10) to obtain that syllable’s frequency. For each word that appeared in the fMRI tasks, we determined the mean, minimum and maximum syllable frequency found in a word.

### 2.4. fMRI analyses

We used SPM12 (Ashburner et al., 2014) in [MATLAB R2022b](https://www.mathworks.com/products/new_products/release2022b.html) (The MathWorks Company) for all fMRI analyses. Negative Zipf word frequency was mean-centered and treated as a parametric modulator of the main effect of word onset within a general linear model (GLM), based on previous findings that increasing word frequency is associated with decreasing brain activation in the VWFA (Kronbichler et al., 2004).

For the reading task, word onset times were provided as part of the dataset. For the speech listening task, we used the Montreal ForcedAligner to generate word onset times from the provided stimulus audio files (McAuliffe et al., 2017). AlignTool was used to force-align auditory task stimuli and determine word onset and offset times (Schillingmann et al., 2018). Matching written transcripts were available within the MOUS dataset for 900 of the auditory sound files, and the remaining 8 audio files were manually transcribed. The AlignTool function ‘Extract On/Offsets’ returned automated onset and offset times, which were manually investigated for accuracy. No corrections of these times were required.

Motion covariates (rotation and translation along each axis) were used as nuisance regressors. When examining correlations between brain activity and word frequency during reading, we controlled for word length (in number of letters) by including it as a parametric regressor of no interest (i.e., with a weighting of zero in the first-level contrast). When examining correlations between brain activity and word frequency during speech listening, we similarly included a control for word duration in milliseconds. Each contrast was aimed at identifying a correlation between brain activity and a specific lexical or sublexical parameter. We thus refer to them as ‘correlation contrasts’ (word frequency during reading, word frequency during speech listening, bigram frequency during reading, and syllable frequency during speech listening).

We used the same parametric regression technique to explore the effect of bigram and syllable frequency. We used the same onset times (corresponding to those of word presentations), but instead of using Zipf word frequency, which reflects the frequency of the word in the corpus, we tested the mean, maximum (Fig. S2, S3), or minimum (Fig. 2, 5) bigram or syllable frequency found in a word as a parametric modulator in separate contrasts of brain activity during reading or speech listening, respectively.

### 2.5. Juxtaposition analyses

Unthresholded statistical parametric t-maps were converted into z-maps using MATLAB’s tcdf function. For each voxel, the cumulative probability of the observed t statistic was computed under the Student’s t-distribution with the residual degrees of freedom estimated from the design matrix. This probability was then mapped to the equivalent standard normal deviate by applying the inverse cumulative distribution function of the standard normal distribution (norminv) using the following equation:

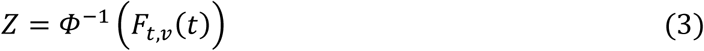

where *t* is the voxelwise t statistic, *v* the residual degrees of freedom, *F*_*t*,*v*_ the t-distribution CDF (computed with tcdf), and *Φ*^−1^ the inverse normal CDF. This transformation preserves the sign of the original t statistic while expressing voxel intensities on the standard normal (Z) scale, allowing different correlation contrasts to be juxtaposed on the same template.

## 3. Results

### 3.1. Lexical frequency regressor reveals significant correlation in the VWFA region and IFG for written words, validating the re-analysis approach

The inverse relationship between VWFA activation in response to written words and word frequency has been established by several prior studies (Graves et al., 2010; Kronbichler et al., 2004; Woolnough et al., 2021). We thus chose to validate our re-analysis approach of the MOUS dataset by first applying it to the scans involving visually presented words. Specifically, we predicted that, in line with its status as a visual lexicon, responses in the VWFA region as well as the downstream inferior frontal gyrus (IFG) would show inverse correlations with word frequency.

Regarding the first prediction, results (Figure 1 and Table 1) revealed negative correlations of responses to written words and lexical frequency in fusiform cortex. Validating the re-analysis approach, the centroid of the peak word frequency correlation cluster (MNI: -42, -53, -13) was localized near the literature coordinates of the VWFA (MNI: -45, -57, -12) (Vogel et al., 2012). In addition, there was a secondary peak in the IFG region, located downstream from the VWFA in the ventral reading stream. Word frequency effects have been previously reported in the IFG (Fiez et al., 1999; Grande et al., 2011; Kronbichler et al., 2004). Interestingly, a third major cluster was found in the left middle temporal gyrus (MTG). This has been less of a focus in the literature, although several studies support the role of this region in reading (Lindenberg & Scheef, 2007; Shaywitz et al., 2004). Finally, we identified a posterior occipital cluster (MNI: -30, -93, -8), in agreement with earlier studies of word selectivity in the brain (Glezer et al., 2015). Supporting theories of the left-lateralization of the reading pathway, no significant clusters were found in the right hemisphere (see Fig. S5 for unthresholded results). Probing for positive correlations with word frequency, only a single cluster was identified in the posterior cingulate cortex (Fig. S1).

**Figure 1.**
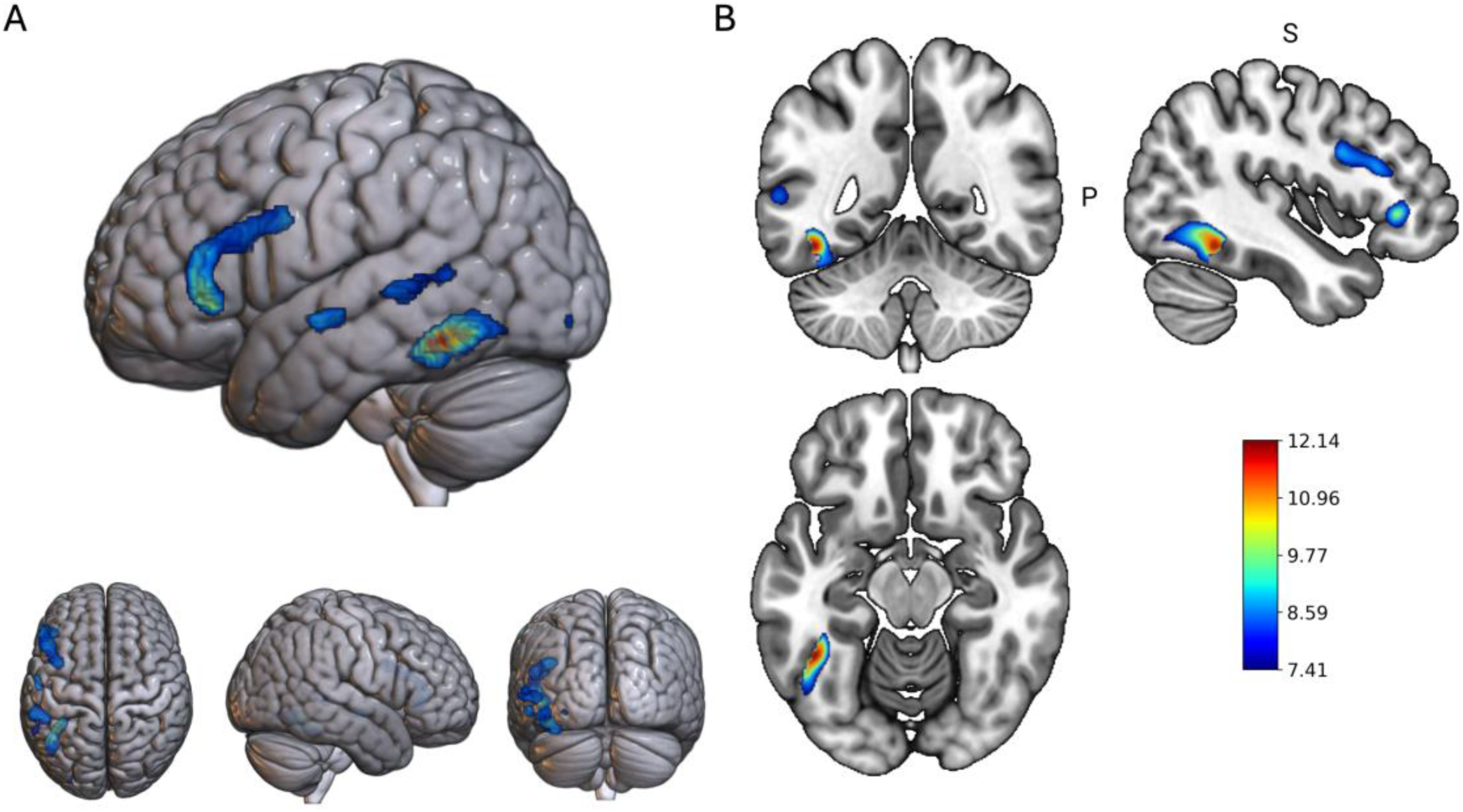
Significant clusters of brain activity negatively correlated with Zipf word frequency during reading. N=99 subjects. Colormap reflects voxel-level T-statistic values. Color bar range spans the height threshold of the t-map at pFWE < 1e⁻⁶ to the t-value of the peak voxel. Extent threshold (kE) = 20 voxels. **A)** 3D-rendering of group-level activation map in MNI space. **B)** Coronal, sagittal, and axial views of the peak cluster (MNI: -44, -50, -14). Renderings were produced using MRIcroGL (Chris & Matthew, 2000).

**Table 1.** Significant clusters showing negative correlation with word frequency during reading. pFWE < 1e⁻⁶, extent threshold of 20 voxels. The table lists the significant clusters of negative correlations with word frequency shown in Figure 1. Cluster centroids were calculated by taking the weighted average of the voxel-wise T-statistics within a surviving cluster in the thresholded T-map (Figure 1). In particular, the strongest correlation (first row) is found close to the literature coordinates of the VWFA.

| Cluster Centroid (MNI) | Region (Brodmann Area) | Extent (voxels) | Cluster peak T-statistic |
| --- | --- | --- | --- |
| -42, -54, -13 | Left fusiform (37) | 444 | 12.14 |
| -48, 24, 15 | Left inferior frontal gyrus, pars triangularis (45); Primary motor cortex (4) | 761 | 10.6 |
| -56, -10, -7 | Left middle temporal gyrus (21) | 94 | 9.36 |
| -56, -40, 4 | Left middle temporal gyrus (21) | 278 | 8.99 |
| -30, -93, -8 | Left occipital pole (18) | 21 | 8.47 |

### 3.2. Correlations with bigram frequency reveal evidence for a hierarchical orthographic representation in the ventral reading stream

We next tested whether our re-analysis approach could identify sublexical representations in the ventral reading stream. Specifically, we tested the hypothesis that regions posterior to the VWFA, such as the OWFA (Strother et al., 2016) a region with sublexical selectivity that lies earlier in the orthographic processing hierarchy, would show inverse correlations between activity and bigram frequency. Given that the VWFA’s responses correlate inversely with lexical frequency (*i.e.,* less frequent words cause stronger responses than more frequent words), we hypothesized that the *least frequent bigram* in a word should dominate the response in sublexical areas. That is, responses in areas posterior to the VWFA should negatively correlate with the frequency of the least frequent bigram (“minimum bigram frequency”) in a word.

Indeed, the minimum bigram frequency correlation contrast (Figure 2) revealed peaks posterior to the VWFA in the ventral occipital-temporal cortex (vOT), earlier in the ventral visual stream’s processing hierarchy (Bruno et al., 2008; Vinckier et al., 2007; Woollams et al., 2011). Notably, in contrast to the word frequency correlation contrast, which only identified significant clusters in the left hemisphere, the bigram frequency correlation contrast identified bilateral occipital clusters.

**Figure 2.**
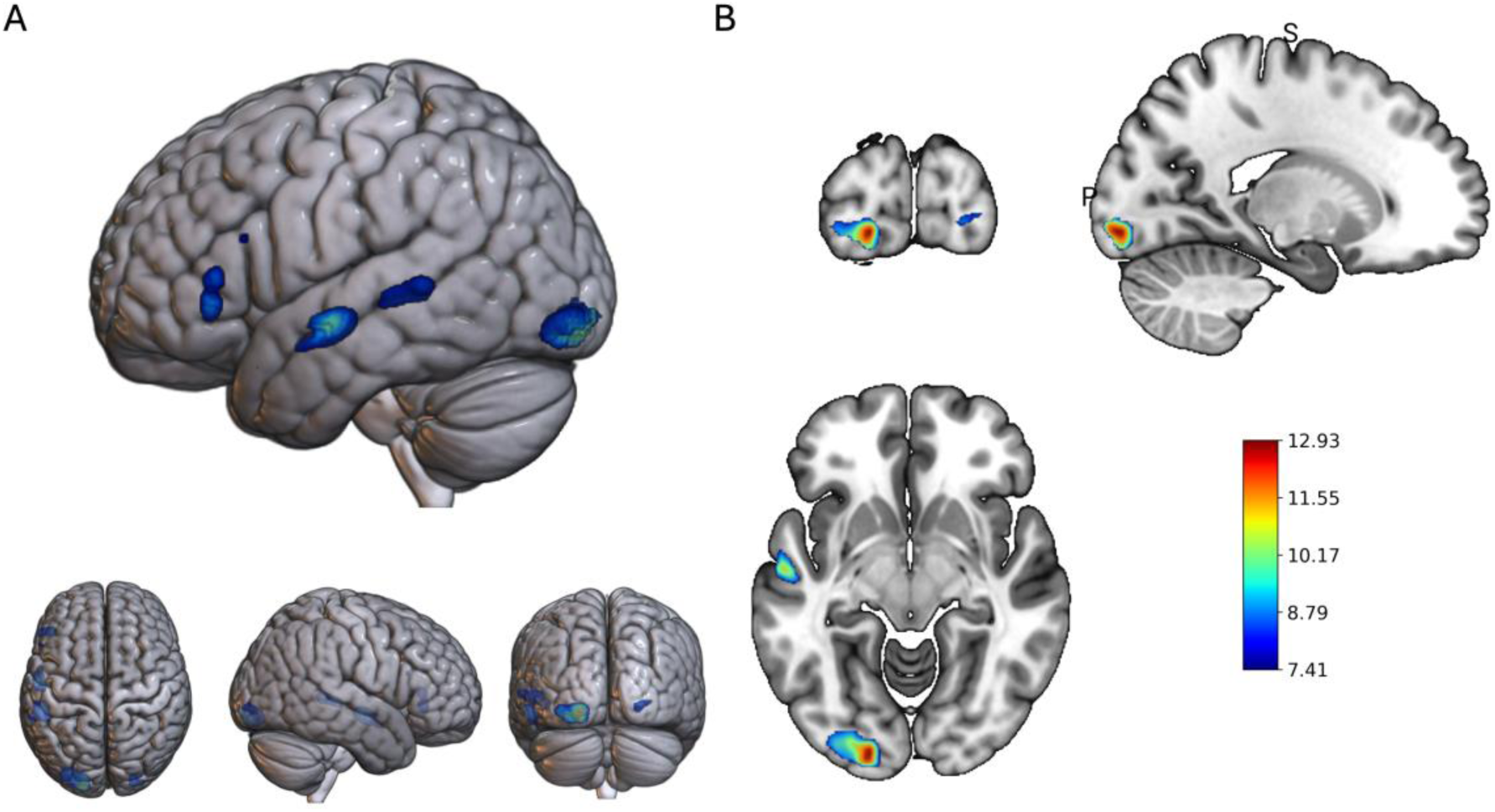
Brain activity negatively correlated with minimum bigram frequency during reading. N = 99 subjects. Colormap reflects voxel-level t-statistic values. Color bar range spans the height threshold of the t-map at pFWE < 1e⁻⁶ to the t-value of the peak voxel. Whole brain rendering shown at right. Extent threshold (kE) = 20 voxels. **A)** 3D-rendering of group-level activation map in MNI space. **B)** Coronal, sagittal, and axial views of the peak cluster, (MNI: -18, -94, -8).

**Table 2.**
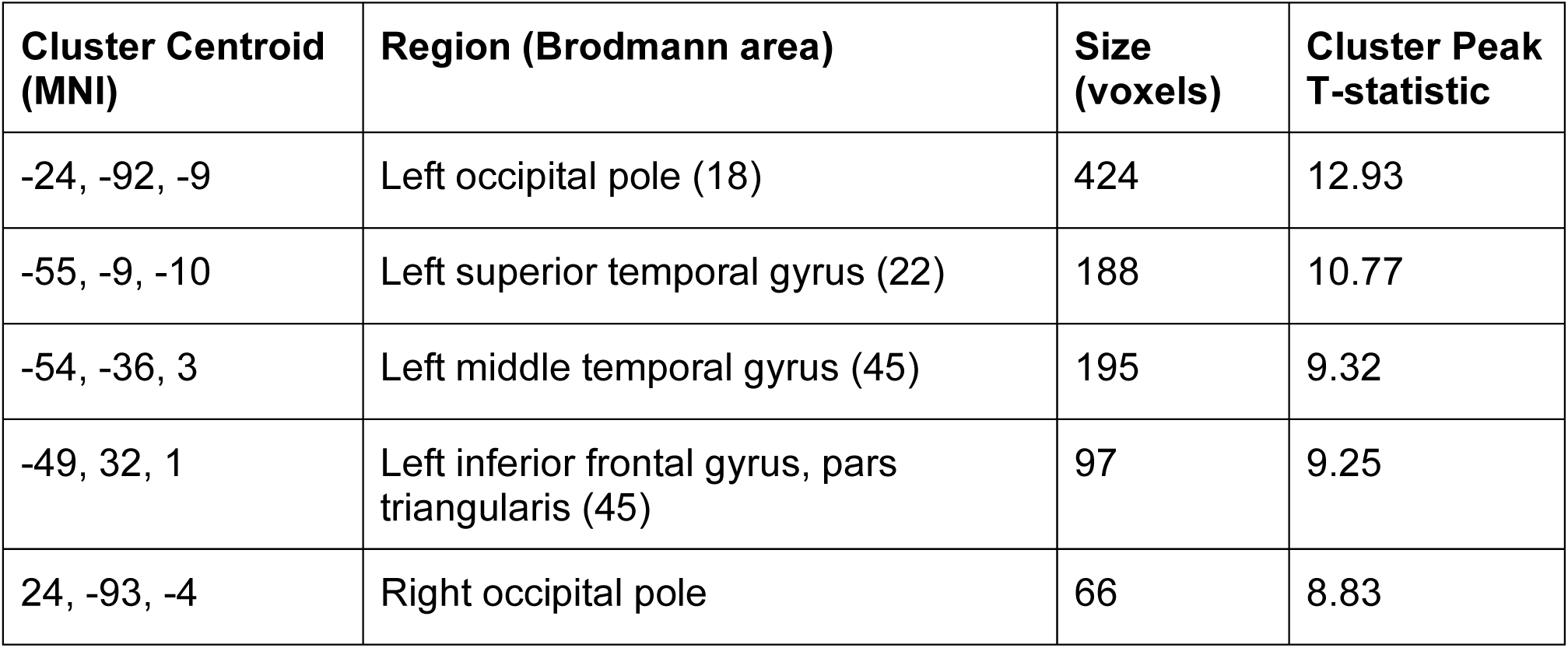
Significant clusters showing negative correlation with minimum bigram frequency contrast during reading. pFWE < 1e⁻⁶, extent threshold of 20 voxels.Cluster centroids were calculated by taking the weighted average of the voxel-wise T-statistics within a surviving cluster in the thresholded T-map (Figure 2).

| Cluster Centroid (MNI) | Region (Brodmann area) | Size (voxels) | Cluster Peak T-statistic |
| --- | --- | --- | --- |
| -24, -92, -9 | Left occipital pole (18) | 424 | 12.93 |
| -55, -9, -10 | Left superior temporal gyrus (22) | 188 | 10.77 |
| -54, -36, 3 | Left middle temporal gyrus (45) | 195 | 9.32 |
| -49, 32, 1 | Left inferior frontal gyrus, pars triangularis (45) | 97 | 9.25 |
| 24, -93, -4 | Right occipital pole | 66 | 8.83 |

**Figure 3.**
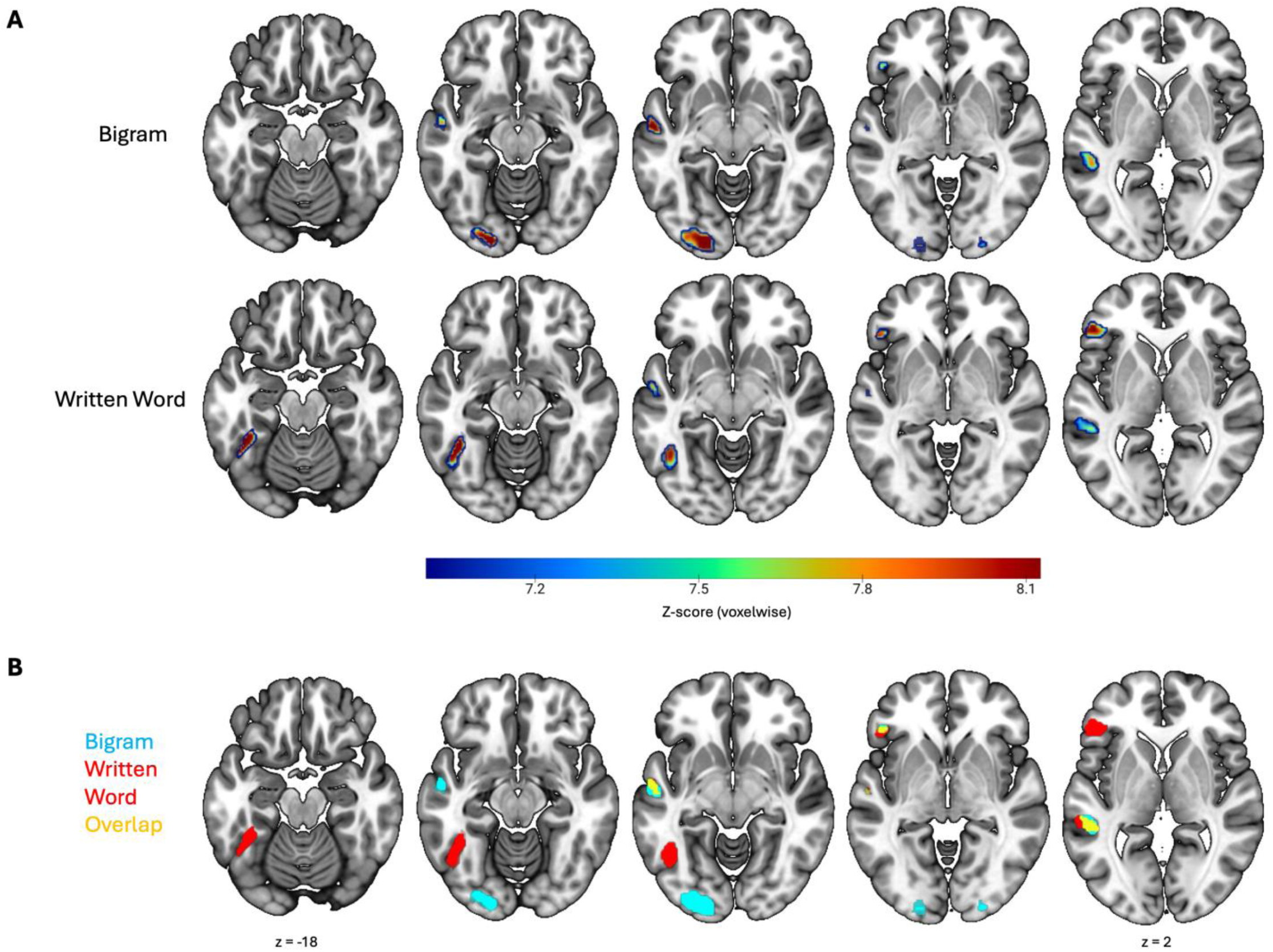
Posterior-to-anterior arrangement of sublexical and lexical selectivity during reading. Axial slices starting at z = -18 are spaced consecutively 5mm apart. **A)** A z-score was derived from the t-statistic of each voxel in the correlation contrast maps of minimum bigram frequency correlation (top row) and word frequency (middle row). **B)** The z-maps in the first two rows were binarized (bigrams in blue, words in red) and juxtaposed, with their spatial overlap shown in yellow.

Interestingly, as in the lexical analysis, we observed some significant correlations in the left IFG and spanning the middle temporal gyrus to superior temporal sulcus. While this could point to sublexical representations in these areas, an alternative explanation is that collinearity between minimum bigram frequency and word frequency (see Fig. S7) captures IFG activation during single-word reading. No significant clusters were found that showed a correlation between the negative maximum or mean bigram frequencies in a word (Fig. S2).

### 3.3. Re-analysis of auditory scans supports the existence of auditory lexical representations in the anteroventral auditory stream

Having validated our re-analysis approach on scans recording brain activation associated with reading printed words, we next turned to the analyses of the auditory scans to probe lexical and sublexical representations for spoken words. As noted in the Introduction, the lexical and sublexical representations of auditory words are much less well understood. Recent work (Damera et al., 2023) has provided evidence for an auditory lexicon in the so-called “Auditory Word Form Area” (AWFA) in the mid-anterior superior temporal cortex, confirming earlier predictions based on a meta-analysis (DeWitt & Rauschecker, 2012)

The contrast examining parametric effect of word frequency during speech listening (Figure 4, unthresholded results in Fig. S6) revealed a group-level cluster in the left mid-anterior superior temporal gyrus (aSTG) with the centroid at (MNI: -52, 3, -16) overlapping with the average location of the AWFA (MNI: −62, −14, 2) identified in our previous study (Damera et al 2023). In addition, significant clusters were found in left parietal and frontal cortex, in line with models of speech processing in human cortex (Hickok & Poeppel, 2007; Rauschecker & Scott, 2009). In particular, there was a cluster in the left IFG that overlapped with the left IFG cluster identified in the lexical correlation analyses for *written* words (see Figure 1 and Table 1), compatible with theories that have posited the left IFG to contain the convergence point of written and spoken word processing hierarchies (Pugh et al., 2001). The analyses also identified a significant cluster in the left angular gyrus, very close to area Spt (Hickok et al., 2003a) thought to play a key role in sensorimotor transformation of speech (Hickok & Poeppel, 2007). Additionally notable, a significant cluster was also found in fusiform cortex, in line with reports of responses to auditory words in the vicinity of the VWFA (Dehaene et al., 2010; Ludersdorfer et al., 2013; Planton et al., 2019; Price & Devlin, 2003; Qin et al., 2021; Yoncheva et al., 2010).

**Figure 4.**
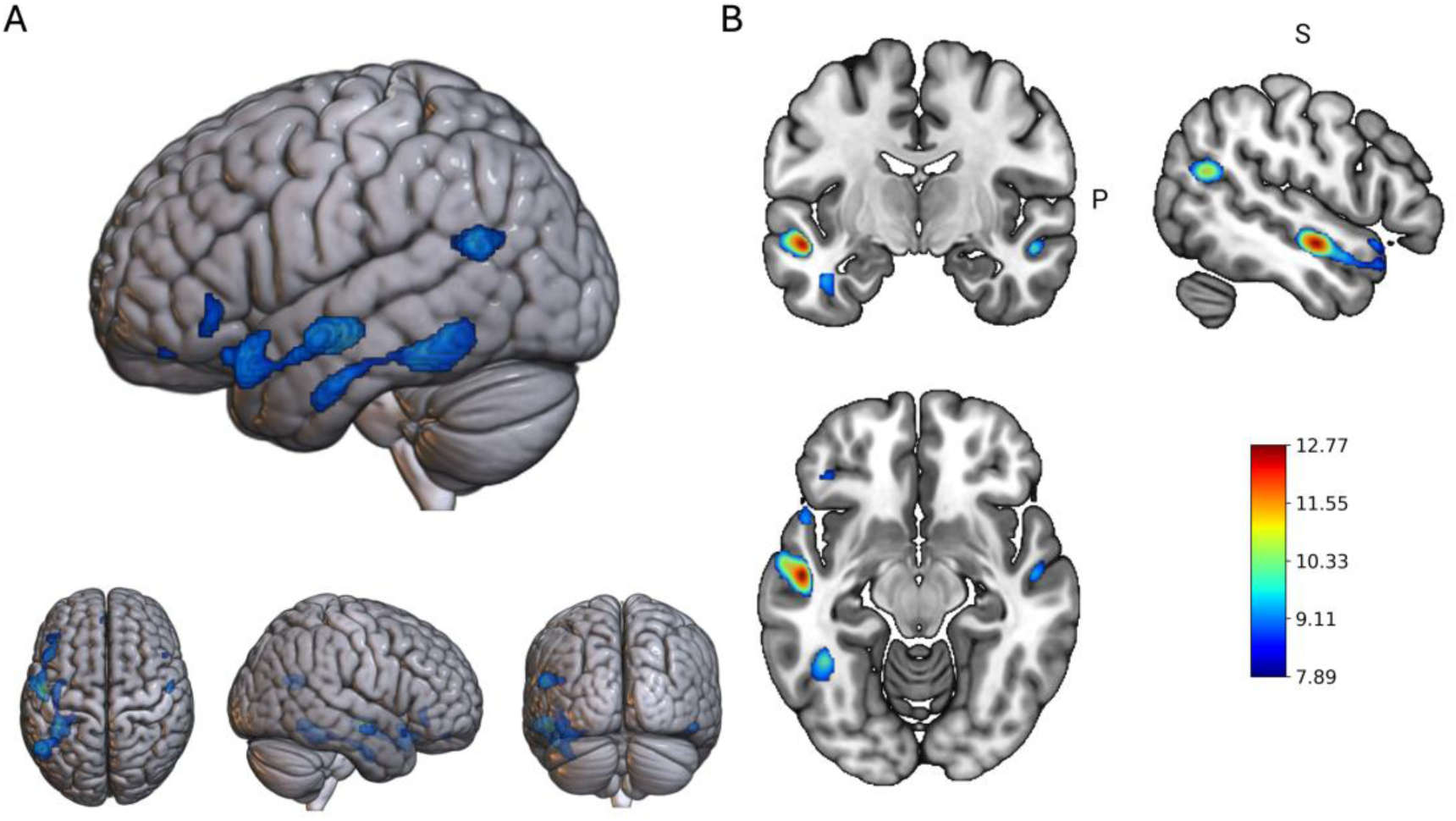
Group-level brain activity negatively correlated with Zipf word frequency during speech listening. N = 101 subjects. Colormap reflects voxel-level T-statistic values. Color bar range spans the height threshold of the t-map at pFWE < 1e⁻⁶ and the t-value of the peak voxel. Extent threshold (kE) = 20 voxels. **A)** 3D-rendering of group-level activation map in MNI space. **B)** Coronal, sagittal, and axial views of the peak cluster, (MNI: -52, -10, -12).

**Table 3.** Significant clusters showing negative correlation with word frequency during speech listening. pFWE < 1e⁻⁶, extent threshold > 20 voxels. The first, most significant ROI denotes the region proximal to the previously described location of the AWFA. Cluster centroids were calculated by taking the weighted average of the voxel-wise T-statistics within a surviving cluster in the thresholded T-map (Figure 4).

| Cluster Centroid (MNI) | Region (Brodmann Area) | Size (voxels) | Cluster Peak T-statistic |
| --- | --- | --- | --- |
| -52, 3, -16 | Left temporal pole (38) | 746 | 12.77 |
| -52, -59, 20 | Left angular gyrus (39) | 317 | 11.04 |
| -39, -36, -20 | Left fusiform gyrus (37) | 751 | 10.61 |
| 53, -8, -15 | Right superior temporal sulcus (21) | 86 | 9.99 |
| -45, 32, -6 | Left anterior superior temporal gyrus (22) | 128 | 9.23 |
| 48, 18, -18 | Right temporal pole | 40 | 8.84 |
| -4, 47, -18 | Left dorsolateral prefrontal cortex | 26 | 8.73 |

Interestingly, there was a difference in the lateralization of these clusters: While significant clusters were found bilaterally in antero-ventral/temporal areas, significant clusters in parietal and frontal areas were found to be exclusively left-lateralized. This intriguing lateralization difference is in line with models of speech processing that have posited bilateral processing of speech in temporal cortex but left-lateralized processing in parietal and frontal cortex (Hickok & Poeppel, 2007). We observed positive correlations with word frequency during speech listening in the right and left superior temporal gyri (Fig. S1).

### 3.4. Evidence for sublexical auditory speech representations in the anteroventral auditory stream supports the existence of an auditory speech processing hierarchy in the auditory “what” stream

In the auditory syllable frequency correlation contrast (Fig. 5, unthresholded results in Fig. S6), we find a cluster in the left mid-superior temporal gyrus (MNI: -61, -17, 1), superior and posterior to the main cluster observed in the word frequency contrast (MNI: -52, 3, -16). This location corresponds to bilateral sublexical phonetic processing centers identified by a prominent meta-analysis (Turkeltaub & Coslett, 2010). Notably, as in the auditory lexical case, a similar cluster was identified in the right hemisphere. Several additional regions with significant activation correlations were observed in response to negative mean (but not maximum) syllable frequency (Fig. S4).

**Figure 5.**
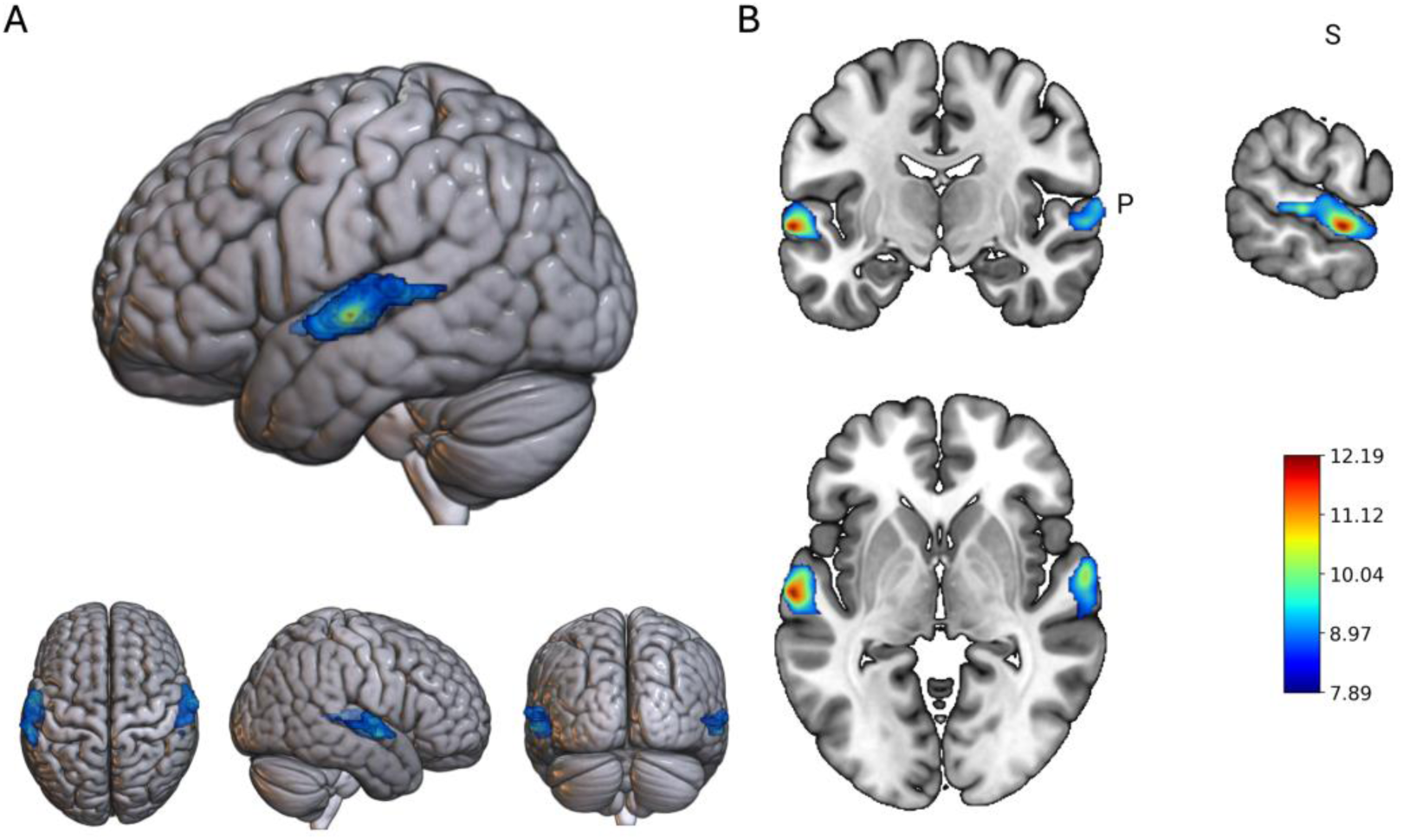
Brain activity negatively correlated with minimum syllable frequency during listening to speech. Color map reflects voxel-level t-statistic values. N = 101 subjects. Color bar range spans the height threshold of the t-map at pFWE< 1e**⁻⁶** and the t-statistic of the peak voxel. Extent threshold (kE) = 20 voxels. **A)** 3D-rendering of group-level activation map in MNI space. **B)** Coronal, sagittal, and axial views of the peak cluster, (MNI: -64, -12, -2).

**Table 4.** Significant clusters showing negative correlation with minimum syllable frequency contrast during speech listening. pFWE < 1e⁻⁶, extent threshold > 20 voxels. This table lists clusters whose activity shows a significant negative correlation with the frequency of the least-frequent syllable in a word. Cluster centroids were calculated by taking the weighted average of the voxel-wise t-statistics within a surviving cluster in the thresholded t-map (Figure 5).

| Cluster Centroid (MNI) | Region (Brodmann Area) | Size (voxels) | Cluster Peak T-statistic |
| --- | --- | --- | --- |
| -61, -17, 1 | Left Superior Temporal Gyrus (22) | 870 | 12.19 |
| 63, -11, 0 | Right Superior Temporal Gyrus (22) | 618 | 11.25 |

**Figure 6.**
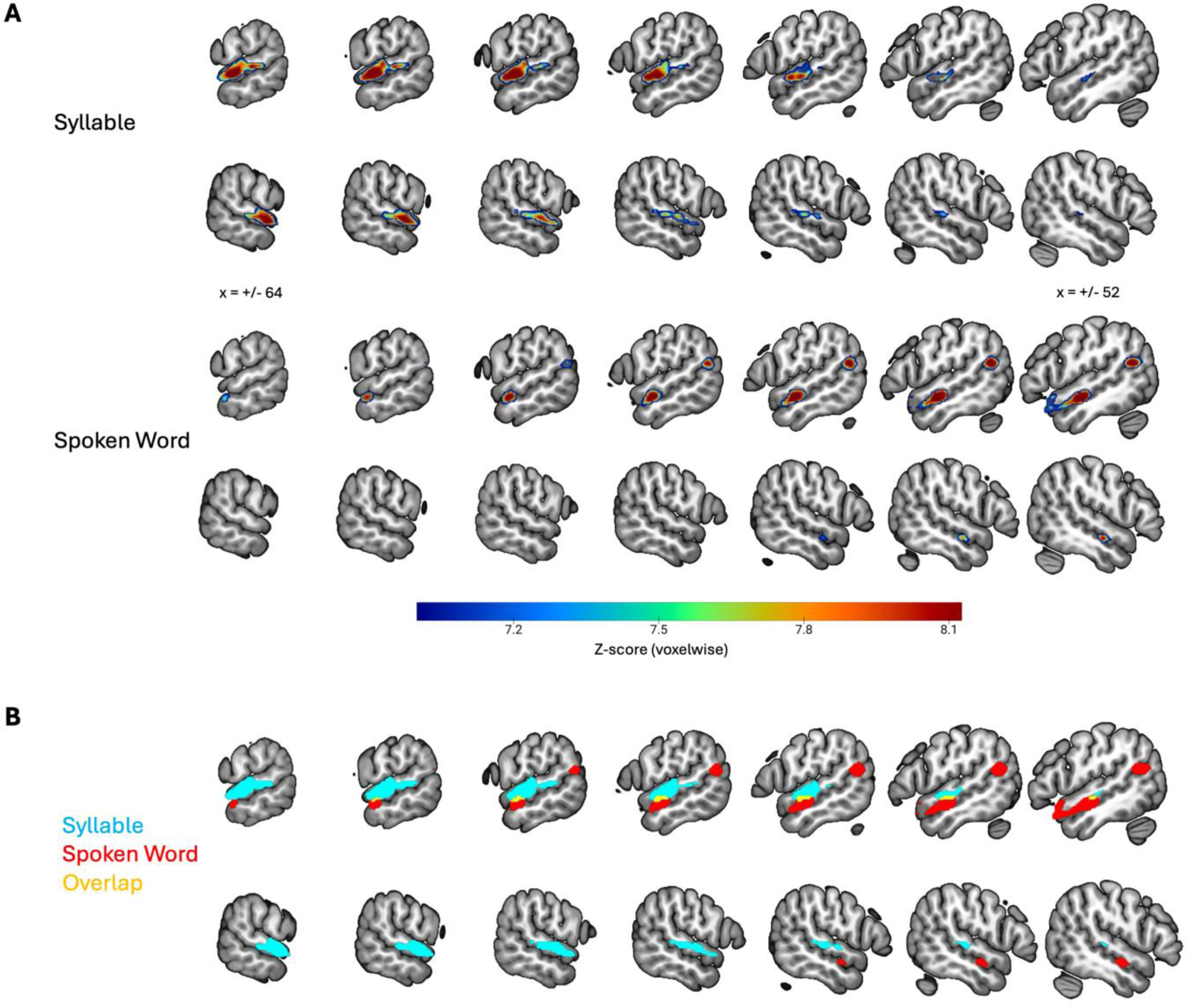
Posterior-to-anterior gradient of lexical selectivity during speech listening in the superior temporal gyrus. Sagittal slices are consecutively spaced 2mm apart. **A)** A z-score was derived from the t-statistic of each voxel in the correlation contrast maps of minimum syllable frequency correlation (top row) and word frequency (middle row). **B)** The z-maps in the first two rows were binarized (syllables in blue, words in red) and juxtaposed, with their spatial overlap shown in yellow.

## 4. Discussion

Learning has been associated with a reduction in the average neuronal response to more familiar stimuli (Anderson et al., 2009; Lim, 2019; Lim et al., 2015; Schulz et al., 2021; Ulanovsky et al., 2003). Linguistic stimuli appear to be no exception: the ‘frequency effect’, wherein rarely-encountered words generate stronger responses than more frequent ones, has been documented for written words across alphabetical and character-based languages (Carreiras et al., 2006; Chee et al., 2003; Graves et al., 2010; Kuo et al., 2003). Previous fMRI studies of the frequency effect during reading (Graves et al., 2010; Kronbichler et al., 2004) that employed parametric designs to explore where in the brain the magnitude of BOLD responses scales with word frequency have identified the VWFA as the site of the strongest (inverse) correlations between activity and word frequency, suggesting that it contains an “orthographic lexicon.” Indeed, subsequent studies using different techniques such as fMRI rapid adaptation (Glezer et al., 2009, 2015) and ECoG (Lochy et al., 2018; Thesen et al., 2012; Woolnough et al., 2021) have provided further confirmatory evidence for the lexicon hypothesis, thus validating the approach of using negative correlations of BOLD response and lexical frequency as a probe for lexical representations in the brain. The frequency effect has previously been shown to also hold for sublexical components of written words, such as bigrams and quadrigrams, thus making it a useful tool for investigating not only lexical, but also sublexical representations of words (Grainger & Whitney, 2004; Graves et al., 2010; Schoonbaert & Grainger, 2004; Vinckier et al., 2007; Woollams et al., 2011).

In the present study, we performed a re-analysis of the MOUS dataset, using the frequency effect to probe for lexical and sublexical representations of written and, for the first time, spoken words in the brain. The MOUS dataset is a multimodal neuroimaging dataset collected from 204 subjects who read and listened to word lists and sentences (Schoffelen et al., 2019). We obtained lexical frequency metrics from the SUBTLEX-NL database, which tracks the occurrences of Dutch words across a corpus consisting of film subtitles, newspapers, books, and other forms of media (Keuleers, Brysbaert, et al., 2010). After segmenting Dutch words into their constituent bigrams and syllables, we calculated frequency metrics for these word parts as well. This enabled us to examine sublexical frequency effects in addition to lexical frequency effects for both written and spoken words

Our study of the reading group confirmed key aspects of the ventral reading pathway and validated our re-analysis approach. The site of the global peak of the correlations between word frequency and activation coincided with the VWFA. An additional cluster was found in the IFG, in agreement with models of reading in the brain (Pugh et al., 2001).

In occipital cortex, our re-analysis identified a main cluster responsive to bigram frequency in the left occipital cortex (posterior to the VWFA), supporting the notion of a simple-to-complex hierarchy for orthographic stimuli in the ventral visual stream. Unlike in our word-frequency correlation contrast, we also found a suprathreshold cluster in the right hemisphere whose activity correlated with bigram frequency. Our main bigram frequency cluster (MNI: -24, -92, -9) corresponds to the literature coordinate of the OWFA (-38,-83,-18), suggesting that bigrams found within whole words that span both hemifields are independently represented by the OWFA, and that experience-dependent tuning (acquired by learning to read) might drive the formation of sublexical, orthographic representations in this region that support the downstream development of lexical representations in the VWFA (Cohen & Dehaene, 2004; Glezer et al., 2015; Strother et al., 2016; Vinckier et al., 2007).

Employing the same parametric regression technique that we validated in our study of the reading group to auditorily presented words, we here report the first neuroimaging analysis of word frequency effects during speech listening. We identified a main cluster in the mid-anterior superior temporal gyrus (MNI: -52, 3, -16) that encompasses and extends anteriorly from the reported location of the Auditory Word Form Area, AWFA (MNI: -62, -14, 2), a possible lexicon for spoken words (Cohen et al., 2004; Damera et al., 2023). This finding supports the role of the auditory anteroventral stream in spoken word recognition and validates the use of the word-frequency correlation method for modeling task-related activations during speech listening.

We also for the first time applied the frequency correlation approach to sublexical auditory processing, focusing on the syllable level. We find a negative linear relationship between the minimum syllable frequency found in a word and BOLD activation in bilateral mid-STG. The left mid-STG cluster (MNI: -61, -17, 1) coincides with the main cluster (MNI: -62, -19, 1) of the activation likelihood map produced by DeWitt & Rauschecker’s 2012 meta-analysis of phoneme repetition suppression studies. Critically, this cluster is posterior to the main cluster identified by the word frequency correlation contrast (MNI: -52, 3, -16), supporting a simple-to-complex (sublexical-to-lexical) hierarchy for processing spoken words that starts in mid-STG and proceeds anteriorly. In this framework, we speculate that the syllable-selective cluster identified by our analyses may contain intermediate acoustic-phonetic representations that are integrated by a word-selective area such as the AWFA. In line with the hypothesis that the strongest activation is associated with the most infrequent words and, by implication, their most infrequent sublexical component, we observed that at the sublexical level, correlations were strongest for the least frequent bigram and syllable, respectively.

Proceeding downstream from the word form areas, across both sensory modalities, the word frequency correlation contrast identified the left IFG. This is consistent with its location at the putative convergence of visual and auditory language perception pathways. The left-lateralization of frontal regions during both reading and speech listening stands in contrast to the more bilateral patterns of selectivity in the temporal lobe, especially for spoken words. We also identified a cluster in the left angular gyrus during speech listening, located close to area Spt in the dorsal speech processing pathway (Hickok et al., 2003b). The auditory dorsal stream computes sensorimotor transformations (Chevillet et al., 2013; Hickok & Poeppel, 2007; Rauschecker, 2011; Rauschecker & Scott, 2009; Rauschecker & Tian, 2000) with some dual-stream models proposing that area Spt is a sensorimotor integration circuit on the basis that it is active during both perception and rehearsal of speech (Hickok et al., 2003a)

Parametric regression techniques like the one used here (that model neural response as a function of stimulus frequency) may be applied to re-analyze other datasets, provided that the frequencies or intensities of stimuli of interest can be quantified and that a sufficient number of stimuli are sampled. Given the rise of ‘naturalistic’ stimulus presentation paradigms (for example, those that employ movie watching) in open-science datasets (Allen et al., 2022), future research efforts may focus on parameterizing ecological variables of interest. For variables that occur with regularity or variable intensity (words, sounds, facial expressions, etc.), parametric regression may be able to localize neural populations selective for them.

Future studies may investigate frequency effects in other languages, such as those with high degrees of homophony (e.g., Japanese), as speakers must access lexical representations from a highly constrained syllabary during speech listening (Tamaoka, 2007). Character-based languages also provide the opportunity to probe word or radical frequency effects during reading. While most studies cited in this manuscript study European languages, which show an inverse relationship between length (in letters or spoken duration) and word frequency, character-based languages show no such (potentially confounding) relationship between orthographic and phonological complexity (Zhang & Xing, 2023).

## 5. Data and Code Availability

Code for all analyses and figures described here and in the Supplement, as well as neuroimaging workflow configuration details, can be found in a Github repository (https://github.com/suneelbanerjee/MOUS_hierarchical-representations). The original dataset is described in Schoffelen et al., 2019 (https://www.nature.com/articles/s41597-019-0020-y).

## 6. Ethics

Study participants whose data were analyzed in this study explicitly consented for the anonymized collected data to be used for research purposes by other researchers. In keeping with the informed consent agreement obtained from the participants and the requirements of both the Ethics Committee and the Radboud University security officer, potentially identifying data (such as imaging data) recorded at the Donders Centre for Cognitive Neuroimaging can only be shared to researchers following explicit approval of a Data Use Agreement. Author S.D. filed this request on behalf of our group.

