## Supplemental Material for "Evidence for hierarchical representations of written and spoken words from an open-science human neuroimaging dataset"

#### Data and Code Availability

Code for all analyses can be found in a Github repository (https://github.com/suneelbanerjee/MOUS_hierarchical-representations). The original dataset is described in Schoffelen et al., 2019 (https://www.nature.com/articles/s41597-019-0020-y)

#### Author Contributions

**Suneel Banerjee**: Conceptualization; Formal analysis; Investigation; Methodology; Project administration; Visualization; Writing – original draft; Writing – reviewing & editing. **Kimberly Jin**: Investigation; Writing – reviewing & editing. **Plamen P. Nikolov**: Investigation; Writing –reviewing & editing. **Phillip Cho**: Investigation; Writing – reviewing & editing. **Vishnu R. Pendri**: Investigation; Writing – reviewing & editing. **Srikanth R. Damera**: Conceptualization; Methodology; Project administration; Writing – reviewing & editing. **Maximilian Riesenhuber**: Conceptualization; Methodology; Project administration; Writing – reviewing & editing.

#### Funding (optional)

National Science Foundation (BCS-1756313) and Georgetown University.

#### Declaration of Competing Interests

The authors declare no competing interests.

#### Acknowledgements

We thank J.M. Schoffelen and colleagues at Radboud University for providing this dataset. Research supported by the National Science Foundation (BCS-1756313) and Georgetown University. This research used resources of the Extreme Science and Engineering Discovery Environment (XSEDE), which was supported by National Science Foundation (1053575).

**Supplementary Material**

SUPPLEMENTAL INFORMATION for *"Evidence for hierarchical representations of written and spoken words from an open-science human neuroimaging dataset."* **Suneel Banerjee, Kimberly Jin, Plamen Nikolov, Philip Cho, Vishnu R. Pendri, Lillian Chang, Srikanth R. Damera, Maximilian Riesenhuber**

### 1. Calculation of bigram and syllable frequencies

#### Filtering of SUBTLEX-NL database

We sourced word frequencies from the "Zipf" column provided in the SUBTLEX-NL lexical database. Before calculating bigram and syllable frequencies for the Dutch language, we cleaned and filtered the SUBTLEX-NL database to remove artifacts left over from text mining of the corpus. These include onomatopoetic words, strings of special characters, misspelled words which do not meaningfully contribute to an estimation of the frequencies of letter combinations and syllables found in everyday Dutch usage.

- **Alphabetic Characters**
  - Retained words composed of standard Dutch letters (including diacritics).
  - Excluded entries containing digits, multiple punctuation marks, or symbols in non-standard positions.
- **Word Length Threshold**
  - Discarded words exceeding a predetermined character limit (25 characters, excluding hyphens). Excessively long strings are likely to be multi-word phrases or processing artifacts rather than individual Dutch words.
  - Discarded single-letter words (e.g., "u") as these could not be parsed into bigrams.
- **Handling 'SPEC' Entries**
  - The tag "SPEC" indicates special categories (e.g., certain proper nouns or abbreviations) in the SUBTLEX-NL database.
  - Retained "SPEC" words only if they were strictly alphabetic. Entries mixing letters with non-standard symbols were excluded.
- **Apostrophe Pattern**
  - Excluded words that begin with a single letter followed by an apostrophe (e.g., "a'trom"), as such patterns are non-standard in Dutch orthography.
- **Repeated Characters and Consecutive Letters**
  - Removed entries composed of a single, repeated letter (e.g., "aaaaa"), as those were likely errors or placeholders rather than genuine words.
  - Excluded any word containing three identical letters in a row (e.g., "aaangepaste").
  - Triple letters are exceedingly rare in standard Dutch and often indicate typos or informal elongations.
- **Text entries incompatible with syllabification algorithm**

- The syllabification algorithm published in Lathouwers et al., 2025 and utilized here to segment Dutch words into syllables is unable to process certain characters (accent marks, apostrophes, umlauts, etc.). Thus, 7605 words in the SUBTLEX-NL database were removed (in addition to the entries excluded by the criteria above) prior to calculating syllable frequency. However, only 4 unique words used in the MOUS language tasks could not be modeled as they contained apostrophes.

- **Number of entries failing each criterion (independently):**

    1) Invalid word pattern: 153807
    2) Exceeding length threshold: 3055
    3) Solely punctuation: 37
    4) SPEC with non-alphabetic chars: 106539
    5) Single-letter + apostrophe: 506
    6) Entirely repeated characters: 173
    7) Triple consecutive letters: 953
    8) Unable to be syllabified: 7605

Of 437503 entries in the database, a total of 159442 were dropped prior to calculating bigram frequencies, and a total of 167047 were dropped prior to calculating syllable frequencies. 29 words used in the MOUS language tasks did not have corresponding entries in the SUBTLEX-NL database; thus the onsets of these words were excluded from all lexical and sublexical Statistical Parametric Mapping (SPM) models.

#### 2. Additional Lexical Contrasts

We also explored potential *positive* correlations between word frequency and brain activity during reading and speech listening.

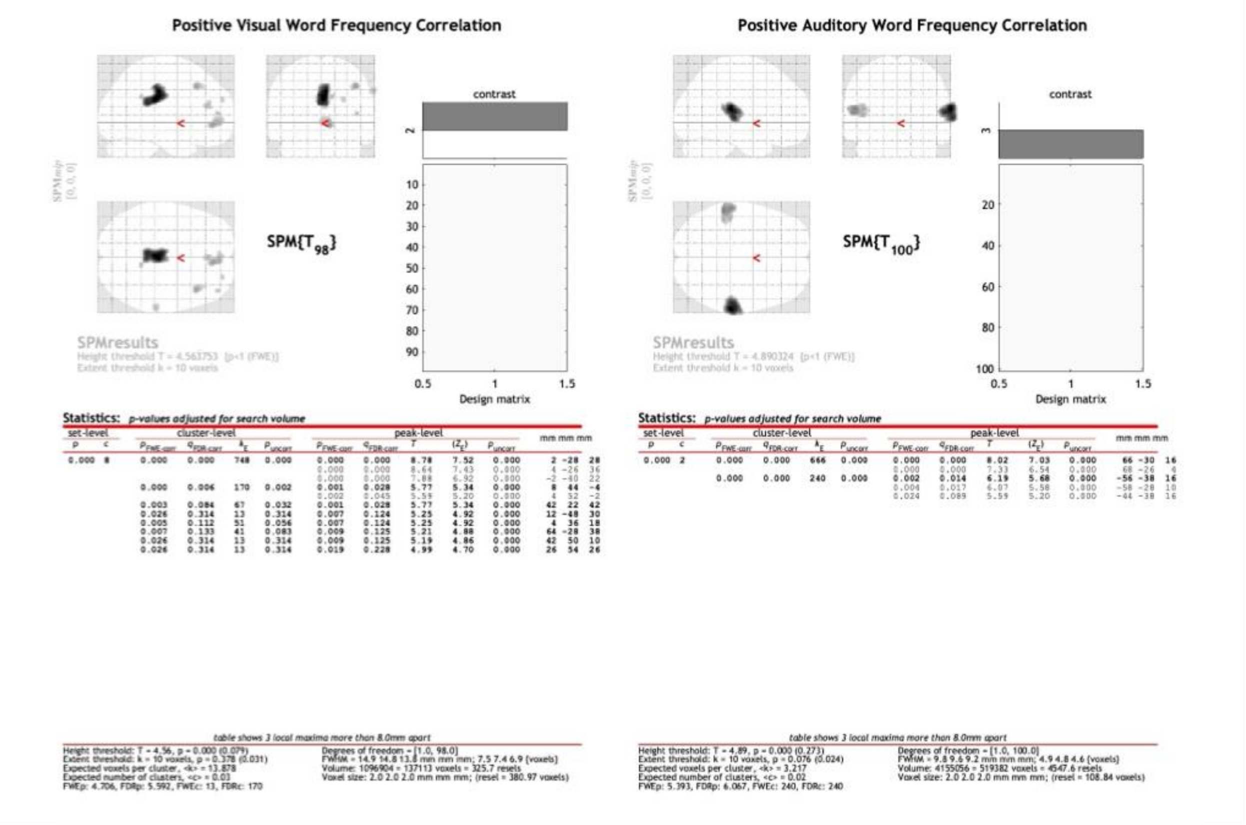

**Figure S1. Positive parametric effects of word frequency during reading (Visual, left) and speech listening (Auditory, right).** A parametric modulator representing Zipf word frequency was tested with a positive contrast weight, indicating an assessment of whether higher word frequency is associated with *increased* BOLD responses. $p < 0.05$ FWE.

#### 3. Additional Sublexical Contrasts

In addition to the negative correlation with minimum bigram/syllable frequencies described in the main text, we also explored positive and negative correlations between the maximum and mean bigram and syllable frequencies found in a word and brain activity during reading and speech listening.

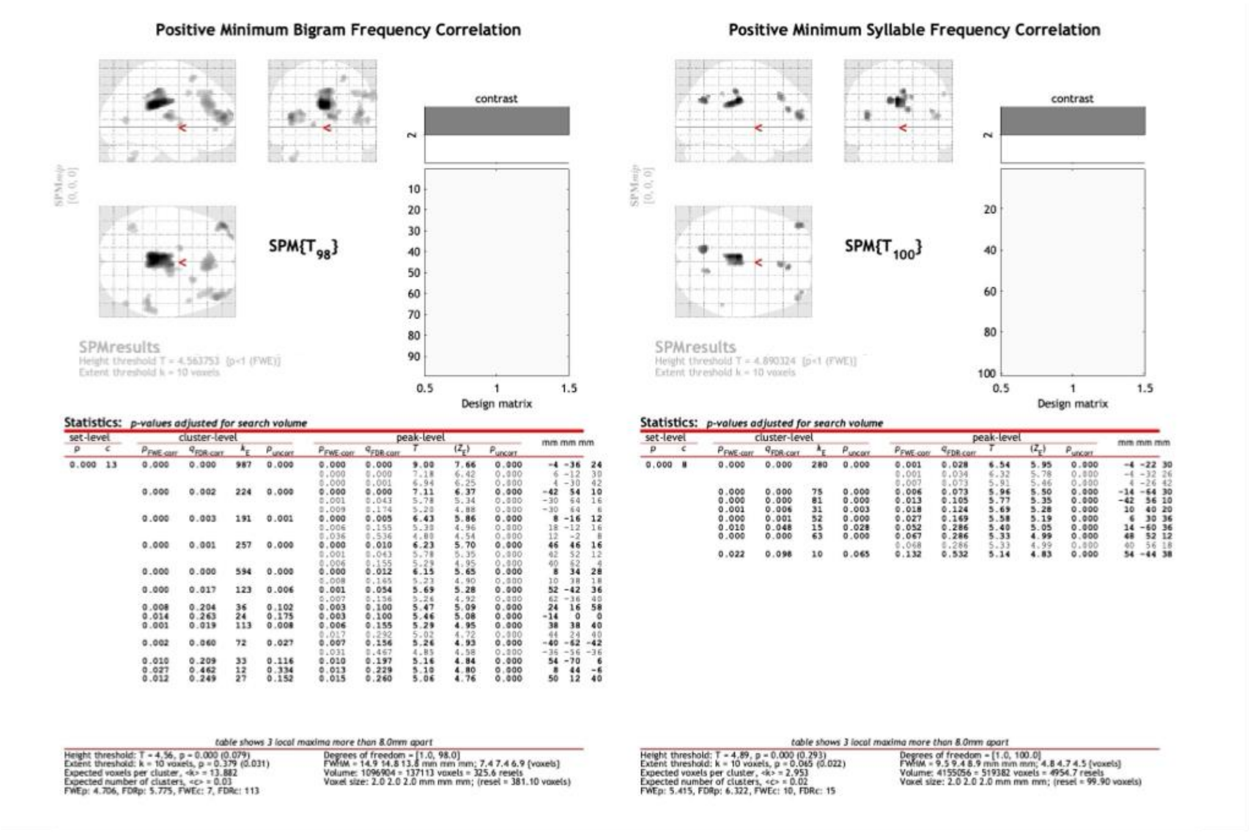


**Figure S2. Positive parametric effects of minimum frequency during reading (left) and minimum syllable frequency during speech listening (right).** A parametric modulator representing Zipf word frequency was tested with a positive contrast weight, indicating an assessment of whether a higher frequency of the rarest sublexical unit is associated with *increased* BOLD responses. pFWE < 1, extent threshold of 10 voxels.

#### Maximum Bigram Frequency (+)

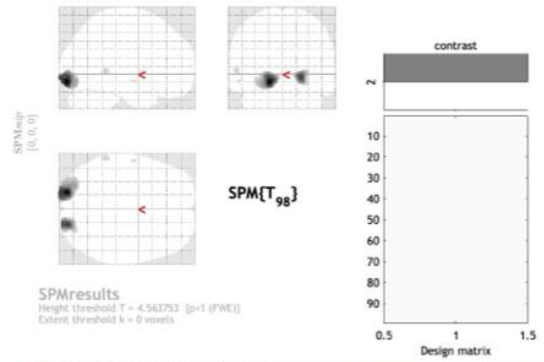


**Statistics:** *p-values adjusted for search volume*

| set-level | | cluster-level | | | | peak-level | | | | | mm mm mm |
|---|---|---|---|---|---|---|---|---|---|---|---|
| $p$ | $c$ | $p_{FWE-corr}$ | $q_{FDR-corr}$ | $k_E$ | $p_{uncorr}$ | $p_{FWE-corr}$ | $q_{FDR-corr}$ | $T$ | $(Z_E)$ | $p_{uncorr}$ | |
| 0.000 | 4 | 0.000 | 0.000 | 718 | 0.000 | 0.000 | 0.000 | 13.67 | Inf | 0.000 | -18 -92 -12 |
| | | | | | | 0.000 | 0.000 | 11.01 | Inf | 0.000 | -18 -100 -8 |
| | | 0.000 | 0.000 | 340 | 0.000 | 0.000 | 0.000 | 10.82 | Inf | 0.000 | 22 -92 -6 |
| | | 0.017 | 0.278 | 20 | 0.209 | 0.011 | 0.158 | 5.16 | 4.84 | 0.000 | -60 -42 2 |
| | | 0.044 | 0.537 | 5 | 0.537 | 0.045 | 0.550 | 4.74 | 4.49 | 0.000 | -56 -8 -10 |

*table shows 3 local maxima more than 8.0mm apart*

Height threshold: T = 4.56, p = 0.000 (0.081)
Extent threshold: k = 0 voxels
Expected voxels per cluster, <k> = 13.558
Expected number of clusters, <c> = 0.08
FWEp: 4.713, FDRp: 10.824, FWEc: 5, FDRc: 340

Degrees of freedom = [1.0, 98.0]
FWHM = 14.7 14.7 13.7 mm mm mm; 7.4 7.4 6.9 {voxels}
Volume: 1096904 = 137113 voxels = 333.4 resels
Voxel size: 2.0 2.0 2.0 mm mm mm; (resel = 372.18 voxels)

#### Maximum Bigram Frequency (-)

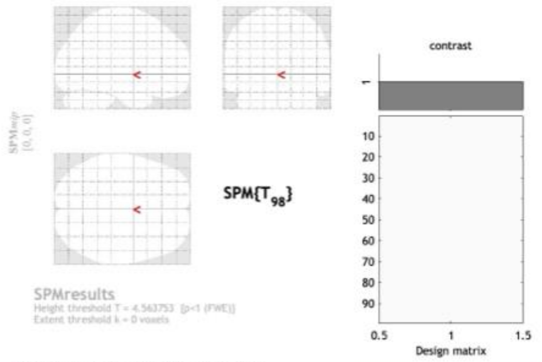


**Statistics:** *p-values adjusted for search volume*

| set-level | | cluster-level | | | | peak-level | | | | | mm mm mm |
|---|---|---|---|---|---|---|---|---|---|---|---|
| $p$ | $c$ | $p_{FWE-corr}$ | $q_{FDR-corr}$ | $k_E$ | $p_{uncorr}$ | $p_{FWE-corr}$ | $q_{FDR-corr}$ | $T$ | $(Z_E)$ | $p_{uncorr}$ | |

*no suprathreshold clusters*

*table shows 3 local maxima more than 8.0mm apart*

Height threshold: T = 4.56, p = 0.000 (0.081)
Extent threshold: k = 0 voxels
Expected voxels per cluster, <k> = 13.558
Expected number of clusters, <c> = 0.08
FWEp: 4.713, FDRp: Inf, FWEc: Inf, FDRc: Inf

Degrees of freedom = [1.0, 98.0]
FWHM = 14.7 14.7 13.7 mm mm mm; 7.4 7.4 6.9 {voxels}
Volume: 1096904 = 137113 voxels = 333.4 resels
Voxel size: 2.0 2.0 2.0 mm mm mm; (resel = 372.18 voxels)

#### Mean Bigram Frequency (+)

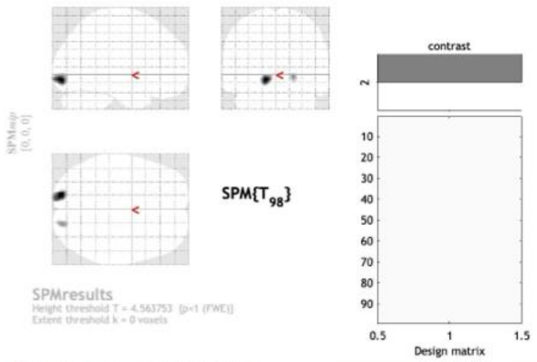


**Statistics:** *p-values adjusted for search volume*

| set-level | | cluster-level | | | | peak-level | | | | | mm mm mm |
|---|---|---|---|---|---|---|---|---|---|---|---|
| $p$ | $c$ | $p_{FWE-corr}$ | $q_{FDR-corr}$ | $k_E$ | $p_{uncorr}$ | $p_{FWE-corr}$ | $q_{FDR-corr}$ | $T$ | $(Z_E)$ | $p_{uncorr}$ | |
| 0.003 | 2 | 0.000 | 0.003 | 174 | 0.002 | 0.000 | 0.000 | 8.22 | 7.15 | 0.000 | -14 -92 -8 |
| | | 0.004 | 0.048 | 56 | 0.048 | 0.001 | 0.010 | 5.81 | 5.37 | 0.000 | 20 -90 -4 |

*table shows 3 local maxima more than 8.0mm apart*

Height threshold: T = 4.56, p = 0.000 (0.078)
Extent threshold: k = 0 voxels
Expected voxels per cluster, <k> = 14.084
Expected number of clusters, <c> = 0.08
FWEp: 4.702, FDRp: 5.812, FWEc: 56, FDRc: 56

Degrees of freedom = [1.0, 98.0]
FWHM = 14.9 14.9 13.9 mm mm mm; 7.5 7.4 6.9 {voxels}
Volume: 1096904 = 137113 voxels = 321.0 resels
Voxel size: 2.0 2.0 2.0 mm mm mm; (resel = 386.62 voxels)

#### Mean Bigram Frequency (-)

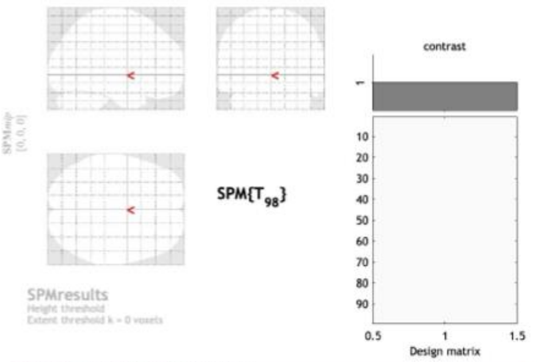


**Statistics:** *p-values adjusted for search volume*

| set-level | | cluster-level | | | | peak-level | | | | | mm mm mm |
|---|---|---|---|---|---|---|---|---|---|---|---|
| $p$ | $c$ | $p_{FWE-corr}$ | $q_{FDR-corr}$ | $k_E$ | $p_{uncorr}$ | $p_{FWE-corr}$ | $q_{FDR-corr}$ | $T$ | $(Z_E)$ | $p_{uncorr}$ | |

*no suprathreshold clusters*

*table shows 3 local maxima more than 8.0mm apart*

Height threshold: T = 4.56, p = 0.000 (0.078)
Extent threshold: k = 0 voxels
Expected voxels per cluster, <k> = 14.084
Expected number of clusters, <c> = 0.08
FWEp: 4.702, FDRp: Inf, FWEc: Inf, FDRc: Inf

Degrees of freedom = [1.0, 98.0]
FWHM = 14.9 14.9 13.9 mm mm mm; 7.5 7.4 6.9 {voxels}
Volume: 1096904 = 137113 voxels = 321.0 resels
Voxel size: 2.0 2.0 2.0 mm mm mm; (resel = 386.62 voxels)

**Figure S3. Parametric effects of maximum (top) and mean (bottom) bigram frequency found in a word during reading.** Parametric modulators for log-transformed maximum and log-transformed mean bigram frequency were tested with a positive (left) and negative (right) contrast weight indicating an assessment of whether higher sublexical unit frequency is associated with reduced or increased BOLD responses, respectively ($p_{FWE} < 1$, extent threshold of 10 voxels).

#### Maximum Syllable Frequency (+)

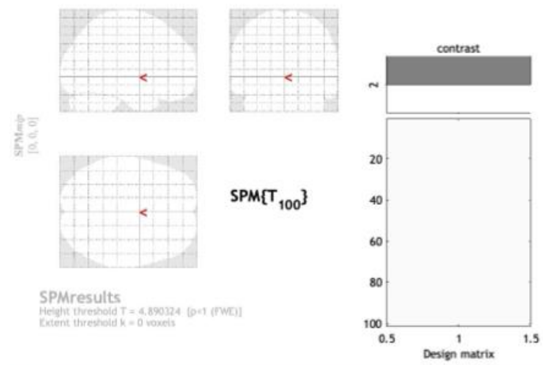


**Statistics:** *p-values adjusted for search volume*

| set-level $p$ | set-level $c$ | cluster-level $p_{FWE-corr}$ | cluster-level $q_{FDR-corr}$ | $k_E$ | cluster-level $p_{uncorr}$ | peak-level $p_{FWE-corr}$ | peak-level $q_{FDR-corr}$ | $T$ | $(Z_E)$ | peak-level $p_{uncorr}$ | mm mm mm |
|---|---|---|---|---|---|---|---|---|---|---|---|

*no suprathreshold clusters*

*table shows 3 local maxima more than 8.0mm apart*

Height threshold: T = 4.89, p = 0.000 (0.302)
Extent threshold: k = 0 voxels
Expected voxels per cluster, <k> = 2.843
Expected number of clusters, <c> = 0.36
FWEp: 5.425, FDRp: Inf, FWEc: Inf, FDRc: Inf

Degrees of freedom = [1.0, 100.0]
FWHM = 9.4 9.3 8.8 mm mm mm; 4.7 4.6 4.4 {voxels}
Volume: 4155056 = 519382 voxels = 5145.4 resels
Voxel size: 2.0 2.0 2.0 mm mm mm; (resel = 96.20 voxels)

#### Maximum Syllable Frequency (-)

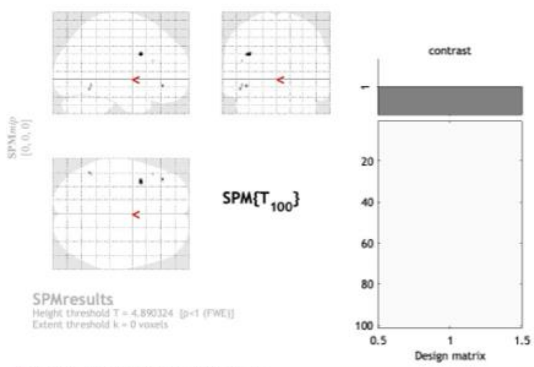


**Statistics:** *p-values adjusted for search volume*

| set-level $p$ | set-level $c$ | cluster-level $p_{FWE-corr}$ | cluster-level $q_{FDR-corr}$ | $k_E$ | cluster-level $p_{uncorr}$ | peak-level $p_{FWE-corr}$ | peak-level $q_{FDR-corr}$ | $T$ | $(Z_E)$ | peak-level $p_{uncorr}$ | mm mm mm |
|---|---|---|---|---|---|---|---|---|---|---|---|
| 0.000 | 5 | 0.015 | 0.213 | 12 | 0.043 | 0.154 | 0.894 | 5.11 | 4.80 | 0.000 | -38 8 28 |
| | | 0.129 | 0.548 | 2 | 0.384 | 0.214 | 0.894 | 5.00 | 4.72 | 0.000 | -40 36 -10 |
| | | 0.179 | 0.548 | 1 | 0.548 | 0.239 | 0.894 | 4.97 | 4.69 | 0.000 | -48 -56 -14 |
| | | 0.129 | 0.548 | 2 | 0.384 | 0.267 | 0.894 | 4.93 | 4.65 | 0.000 | -46 -54 -10 |
| | | 0.179 | 0.548 | 1 | 0.548 | 0.275 | 0.894 | 4.92 | 4.65 | 0.000 | -46 24 20 |

*table shows 3 local maxima more than 8.0mm apart*

Height threshold: T = 4.89, p = 0.000 (0.302)
Extent threshold: k = 0 voxels
Expected voxels per cluster, <k> = 2.843
Expected number of clusters, <c> = 0.36
FWEp: 5.425, FDRp: Inf, FWEc: 12, FDRc: Inf

Degrees of freedom = [1.0, 100.0]
FWHM = 9.4 9.3 8.8 mm mm mm; 4.7 4.6 4.4 {voxels}
Volume: 4155056 = 519382 voxels = 5145.4 resels
Voxel size: 2.0 2.0 2.0 mm mm mm; (resel = 96.20 voxels)

#### Mean Syllable Frequency (+)

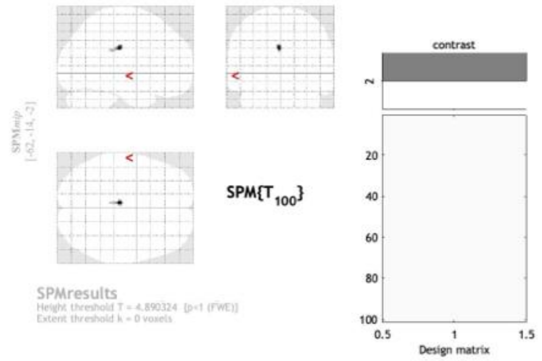


**Statistics:** *p-values adjusted for search volume*

| cluster-level $p_{FWE-corr}$ | cluster-level $q_{FDR-corr}$ | $k_E$ | cluster-level $p_{uncorr}$ | peak-level $p_{FWE-corr}$ | peak-level $q_{FDR-corr}$ | $T$ | $(Z_E)$ | peak-level $p_{uncorr}$ | mm mm mm |
|---|---|---|---|---|---|---|---|---|---|
| 0.003 | 0.009 | 22 | 0.009 | 0.051 | 0.295 | 5.42 | 5.06 | 0.000 | -4 -24 30 |
| | | | | 0.218 | 0.692 | 4.99 | 4.71 | 0.000 | -4 -32 26 |

*table shows 3 local maxima more than 8.0mm apart*

Height threshold: T = 4.89, p = 0.000 (0.299)
Extent threshold: k = 0 voxels
Expected voxels per cluster, <k> = 2.881
Expected number of clusters, <c> = 0.36
FWEp: 5.422, FDRp: Inf, FWEc: 22, FDRc: 22

Degrees of freedom = [1.0, 100.0]
FWHM = 9.5 9.3 8.9 mm mm mm; 4.7 4.7 4.4 {voxels}
Volume: 4155056 = 519382 voxels = 5078.6 resels
Voxel size: 2.0 2.0 2.0 mm mm mm; (resel = 97.46 voxels)

#### Mean Syllable Frequency (-)

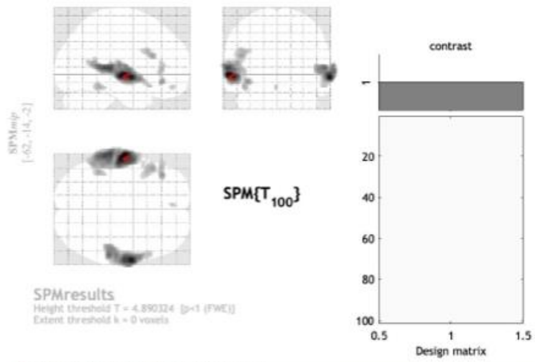


**Statistics:** *p-values adjusted for search volume*

| set-level $p$ | set-level $c$ | cluster-level $p_{FWE-corr}$ | cluster-level $q_{FDR-corr}$ | $k_E$ | cluster-level $p_{uncorr}$ | peak-level $p_{FWE-corr}$ | peak-level $q_{FDR-corr}$ | $T$ | $(Z_E)$ | peak-level $p_{uncorr}$ | mm mm mm |
|---|---|---|---|---|---|---|---|---|---|---|---|
| 0.000 | 7 | 0.000 | 0.000 | 1857 | 0.000 | 0.000 | 0.000 | 9.42 | Inf | 0.000 | -62 -14 -2 |
| | | | | | | 0.000 | 0.000 | 7.86 | 6.92 | 0.000 | -64 -32 6 |
| | | | | | | 0.000 | 0.002 | 6.72 | 6.09 | 0.000 | -62 -20 8 |
| | | 0.000 | 0.000 | 1107 | 0.000 | 0.000 | 0.000 | 8.89 | 7.61 | 0.000 | 64 -4 -6 |
| | | | | | | 0.000 | 0.000 | 8.02 | 7.03 | 0.000 | 64 -12 -2 |
| | | | | | | 0.000 | 0.000 | 7.27 | 6.50 | 0.000 | 58 -32 6 |
| | | 0.000 | 0.000 | 152 | 0.000 | 0.001 | 0.005 | 6.44 | 5.88 | 0.000 | -40 34 -10 |
| | | | | | | 0.008 | 0.033 | 5.89 | 5.44 | 0.000 | -48 34 4 |
| | | | | | | 0.081 | 0.250 | 5.29 | 4.95 | 0.000 | -56 34 12 |
| | | 0.000 | 0.000 | 171 | 0.000 | 0.003 | 0.011 | 6.18 | 5.68 | 0.000 | -48 22 20 |
| | | | | | | 0.010 | 0.038 | 5.83 | 5.40 | 0.000 | -46 16 26 |
| | | 0.000 | 0.000 | 79 | 0.000 | 0.009 | 0.033 | 5.88 | 5.44 | 0.000 | -44 -46 -16 |
| | | 0.007 | 0.023 | 17 | 0.019 | 0.062 | 0.198 | 5.36 | 5.02 | 0.000 | -4 48 -18 |
| | | 0.011 | 0.031 | 14 | 0.031 | 0.137 | 0.414 | 5.14 | 4.83 | 0.000 | -50 10 -10 |

*table shows 3 local maxima more than 8.0mm apart*

Height threshold: T = 4.89, p = 0.000 (0.299)
Extent threshold: k = 0 voxels
Expected voxels per cluster, <k> = 2.881
Expected number of clusters, <c> = 0.36
FWEp: 5.422, FDRp: 5.835, FWEc: 14, FDRc: 14

Degrees of freedom = [1.0, 100.0]
FWHM = 9.5 9.3 8.9 mm mm mm; 4.7 4.7 4.4 {voxels}
Volume: 4155056 = 519382 voxels = 5078.6 resels
Voxel size: 2.0 2.0 2.0 mm mm mm; (resel = 97.46 voxels)

**Figure S4. Parametric effects of maximum and mean syllable frequency found in a word during speech listening.** Parametric modulators for log-transformed maximum and log-transformed mean syllable were tested with a positive (left) or negative (right) contrast weight indicating an assessment of whether higher sublexical unit frequency is associated with increased or reduced BOLD responses, respectively ($p_{FWE} < 1$, extent threshold of 10 voxels).

#### Unthresholded Contrast Images

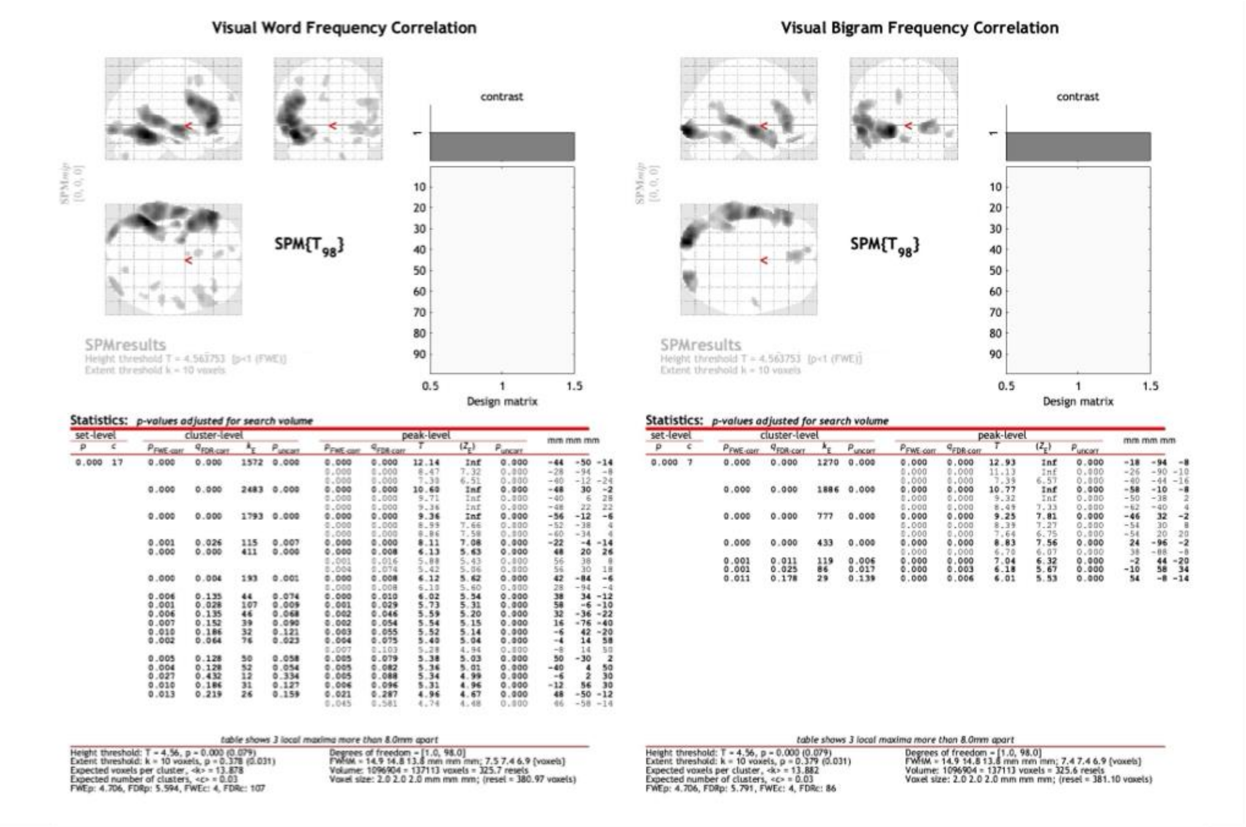


**Figure S5. Unthresholded parametric effects of word frequency and bigram frequency during reading.** pFWE < 1, extent threshold of 10 voxels. These contrasts correspond to those applied in Figures 1-3.

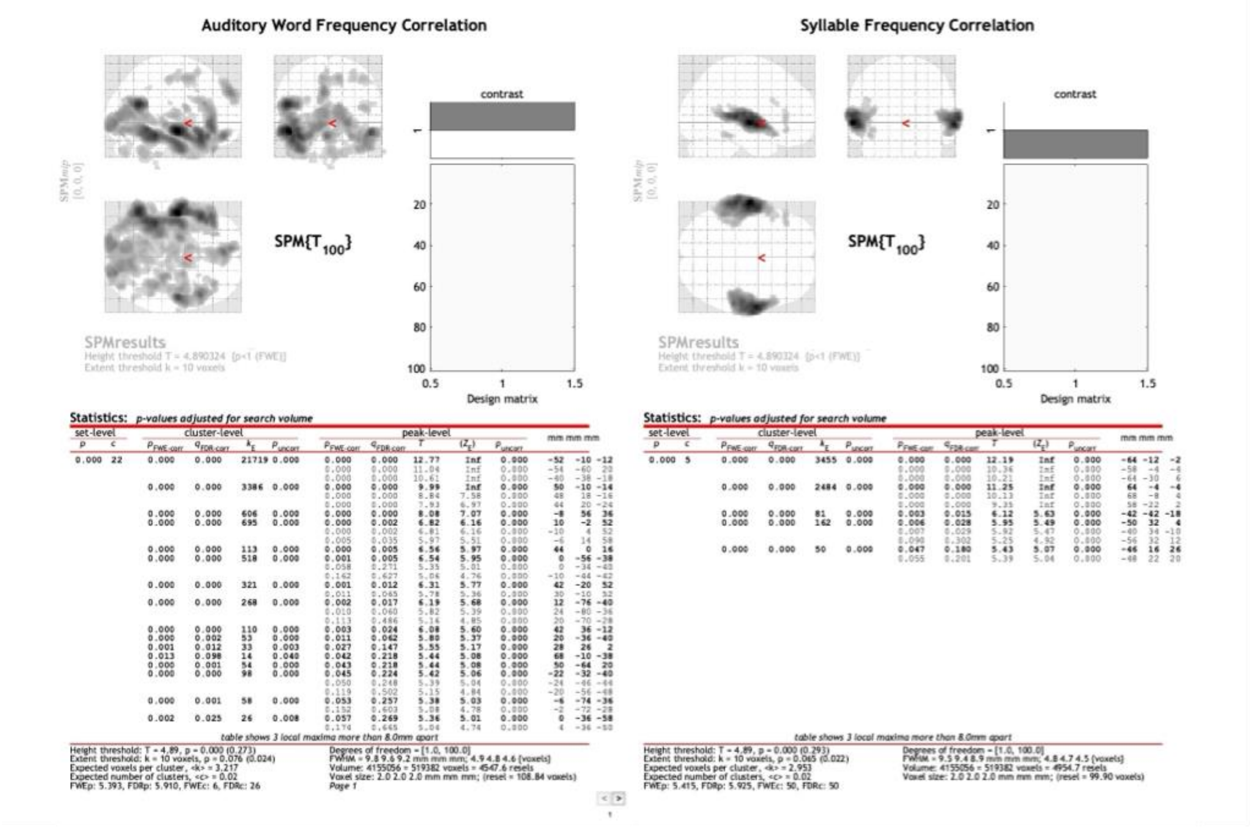


**Figure S6. Unthresholded parametric effects of word frequency and syllable frequency during speech listening.** pFWE < 1, extent threshold of 10 voxels. These contrasts correspond to those applied in Figures 4-6.

### 4. Correlations between lexical and sublexical features

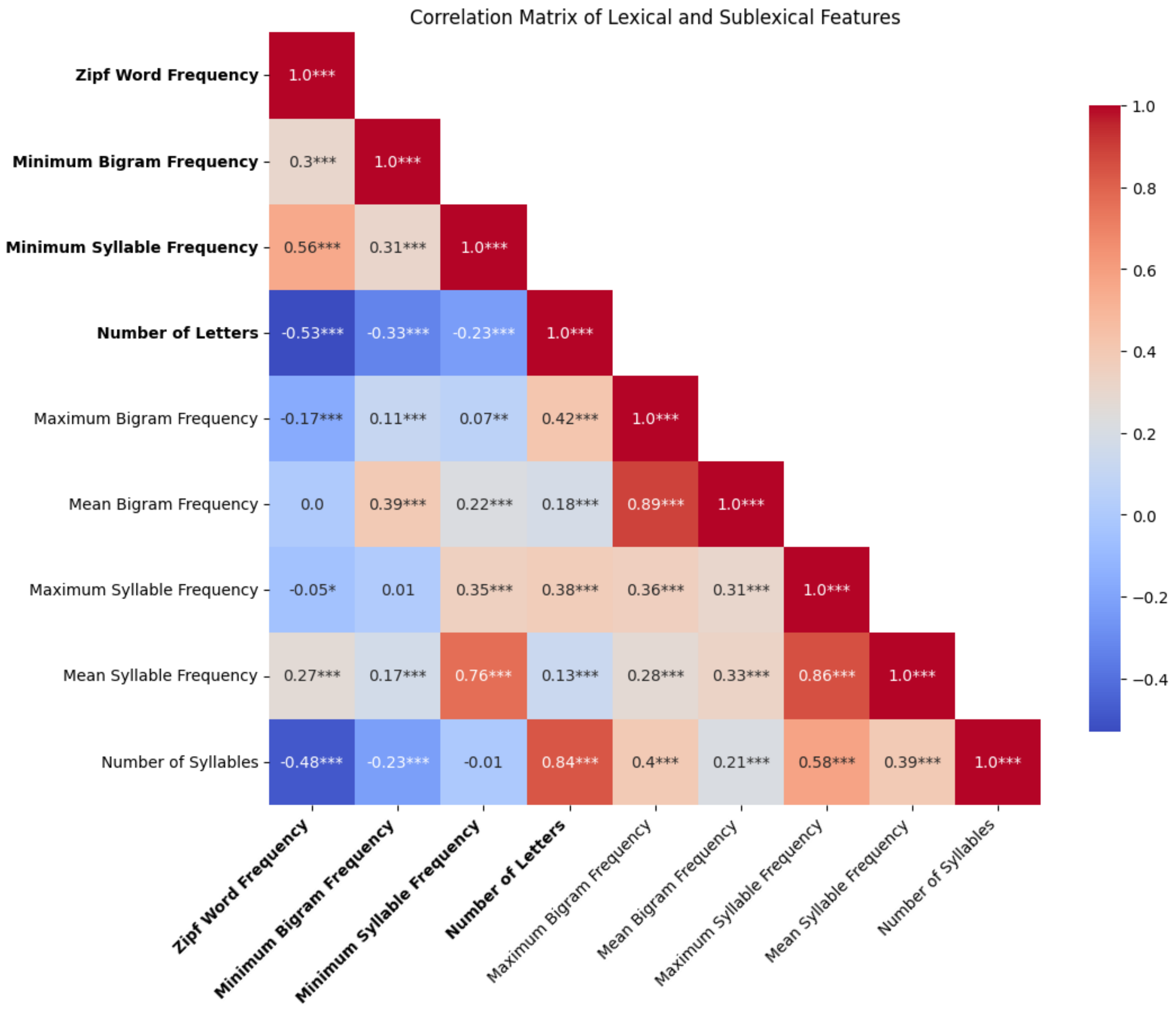


**Figure S7. Correlations between lexical and sublexical features.** The matrix displays pairwise Pearson correlation coefficients among nine word-level parameters, including word frequency, log-transformed bigram and syllable frequency measures, and structural characteristics (letter and syllable counts) for all modeled words (n = 1910) that appeared in the MOUS language tasks. Asterisks denote statistical significance of each correlation coefficient (p < .001: ****). Bolded axis labels indicate the four features that were ultimately selected as parametric modulators in the fMRI general linear models. Not shown above is spoken word duration in seconds, which was included as a parametric modulator in the auditory word frequency correlation contrast, but is not a constant lexical characteristic. While log-transformed maximum and mean frequencies for syllables and bigrams were initially considered as candidate parametric regressors for fMRI analysis they were excluded due to lack of explanatory power (see Figures S3 & S4).

#### 5. Additional Analyses

Our analyses in the paper focused on correlations with lexical and sublexical frequency alone. This not only allows comparison to the prior literature that used the frequency correlation technique (Bruno et al., 2008; Carreiras et al., 2006; Chee et al., 2003; Graves et al., 2010; Kronbichler et al., 2004), but the advantage of these single correlations is their straightforward interpretability, testing the hypotheses that the activation of a lexical representation consisting of neurons holistically tuned to words (see (Glezer et al., 2009) correlates inversely with word frequency, and the activation of a sublexical representation consisting of neurons tuned to sublexical units correlates inversely with the frequency of the most infrequent (and often the first, see Fig R3) sublexical unit.

To explore the questions what activations in response to bigram/syllable frequency are not shared by written/auditory word frequency, and vice versa, we have also performed additional analyses that included sublexical and lexical frequency measures in the same general linear model (GLM) for each sensory modality. In each sensory modality, we explore the effect of one parameter while treating the other as a parameter of no interest. This allows us to examine the effect of word frequency above and beyond that which can be explained by bigram/syllable frequency and vice-versa.

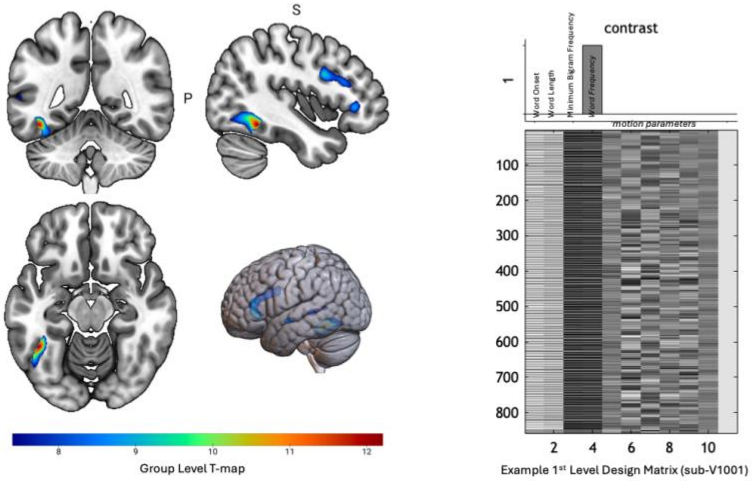


| Cluster Peak (MNI) | Region (Brodmann Area) | Extent (voxels) | Peak T-value |
|---|---|---|---|
| -44, -50, -14 | *Left fusiform gyrus (37)* | 409 | 12.21* |
| -52, 34, 2 | *Left inferior frontal gyrus pars triangularis (45)* | 678 | 10.12 |
| -56, -8, -8 | *Left superior temporal gyrus (22)* | 58 | 8.95 |
| -60, -34, 4 | *Left middle temporal gyrus (21)* | 121 | 8.49 |
| -60, -52, 8 | *Left angular gyrus (11)* | 20 | 7.91 |

**Figure S8. The effect of word frequency when including bigram frequency as a regressor of no interest in a combined model.** Top left: Rendering of group-level effect of word frequency (controlling for word length and minimum bigram frequency). Top right: Example SPM12 contrast vector and design matrix showing all parametric regressors included in a general linear model for a single subject. Bottom: Table listing all significant clusters (pFWE < $1e^{-6}$, extent threshold of 20 voxels).

We find significant clusters associated with word frequency in the VWFA region during reading, even when including minimum bigram frequency as a regressor of no interest. The strength of our peak cluster (asterisk), located in the VWFA, increases by 0.07 (relative to Table 1) when including bigram frequency as a regressor of no interest. The occipital cluster we previously observed not only in the sublexical (Table 2) but also in the word frequency analysis (Table 1) no longer shows significant correlation with word frequency when including minimum bigram frequency as a regressor of no interest. These results reinforce that the VWFA is selectively sensitive to whole-word representations beyond simple letter co-occurrence patterns, and that the posterior cluster is selective at the sublexical but not the lexical level.

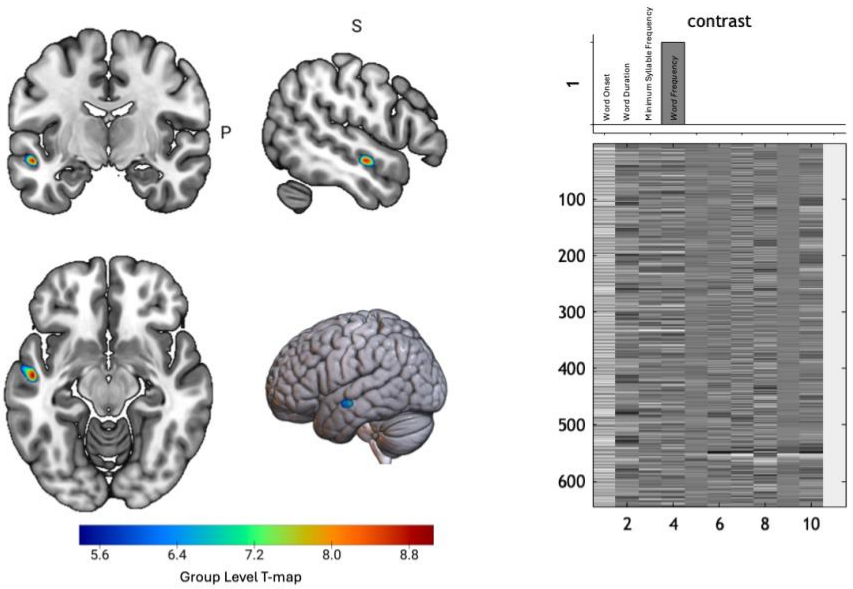


| Coordinates | Region (Brodmann Area) | Extent | Peak T- value |
|---|---|---|---|
| -52, -10, -12 | *Left Superior Temporal Gyrus (22)* | 80 | 9.05 |

**Figure S9. The effect of word frequency when including syllable frequency as a regressor of no interest in a combined model.** Top left: Rendering of group-level effect of word frequency (controlling for word duration and minimum syllable frequency). Top right: Example SPM12 contrast vector and design matrix showing all parametric regressors included in a general linear model for a single subject. Bottom: Table listing all significant clusters (pFWE < $1e^{-6}$, extent threshold of 20 voxels).

For speech listening (Fig. S9 above), we find only one surviving cluster in in the superior temporal gyrus that is associated with word frequency when including syllable frequency as a regressor of no interest. The location of this cluster corresponds to that reported as the AWFA in the main manuscript. Thus, even when controlling for syllable frequency, we still observe that the activity of the AWFA is inversely correlated with the frequency of spoken words, providing further support for the AWFA's hypothesized role as an auditory lexicon.

In contrast to the lexical frequency correlation results, we do not find noteworthy significant correlation clusters associated with sublexical frequency when including lexical frequency in the same model.
